# Evolution and spread of multi-adapted pathogens in a spatially heterogeneous environment

**DOI:** 10.1101/2022.07.16.500289

**Authors:** Quentin Griette, Matthieu Alfaro, Gaël Raoul, Sylvain Gandon

## Abstract

The emergence and the spread of multi-adapted pathogens is often viewed as a slow process resulting from the incremental accumulation of single adaptations. In bacteria, for instance, multidrug resistance to antibiotics may result from the sequential acquisition of single drug resistance to different antibiotics. In phytopathogens, the ability to infect different resistant varieties of crops may also result from the accumulation of distinct virulence genes. Here we use a general epidemiological model to analyse the evolution of pathogen adaptations throughout an epidemic spreading in a heterogeneous host population where selection varies periodically in space. This spatially heterogeneous selection may result from the use of different drugs, different vaccines or different crop varieties in agriculture. We study both the transient evolution of pathogen adaptation at the front of the epidemic and the long-term evolution far behind the epidemic front. We identify five different types of epidemic profiles that may arise from different combinations of spatial heterogeneity and the cost of multi-adaptation. In particular, we show that multi-adaptation can drive epidemic spread, while the evolution of single-adaptation may only occur in a second phase, when the pathogen specializes on local selective pressures. Indeed, a generalist pathogen with multiple adaptations can outpace the spread of a coalition of specialist pathogens when selection varies frequently in space. This result is amplified in finite host populations because demographic stochasticty can lead to the extinction of maladapted pathogens specialised to a local selective pressure. Our work has important implications for the management of multiple drugs and vaccines against pathogens but also for the optimal deployment of resistant varieties in agriculture.

## 1 Introduction

The rise of antimicrobial resistance is a major public-health issue [1]. In particular, the emergence and the spread of bacteria carrying resistance to multiple antibiotics erode our ability to cure these infections [2, 3]. The evolution of multiresistant pathogens is often viewed as the result of the slow accumulation of resistance genes after the persistent use of different drugs in the host population. Similarly, pathogen adaptation to different vaccines or to different resistant varieties of crops may involve the sequential acquisition of distinct escape mutations. Under this scenario, the evolution of a a broader range of resistance to drugs (i.e., multi-adaptation) is viewed as an unavoidable long term consequence of the use of multiple drugs. However, the long-term coexistence of bacteria that differ in the range of drug resistance (i.e., single-adaptation) challenges this view [4, 5]. Several theoretical studies highlighted the importance of different forms of heterogeneities in the structure of the host population to maintain the coexistence between sensitive and resistant bacteria [6, 7, 8, 9, 10, 11]. For instance, spatial variation in drug treatment can readily maintain this long-term coexistence when migration is limited among host populations [6, 7, 9]. Spatial variation in selection may also emerge in spatially spreading epidemics where selection acting at the frontline of the epidemic may differ from selection behind the front [12, 13, 14]. These earlier studies focused on the evolution of pathogen life-history traits like virulence and transmission in epidemics spreading in a homogeneous environment. Here we explore the interplay between pathogen epidemiology and the evolution of drug resistance in a spatially heterogeneous environment.

The spread of epidemics in homogeneous environments can be described by a *travelling front* where the pathogen propagates through space at a constant speed [13, 14, 15]. In a spatially heterogeneous environment, the variation of the quality of the habitat affects the speed of epidemic spread. Yet, if the spatial variation is periodic, the natural extension of the travelling front is the so-called *pulsating front* characterized by its average speed [15, 16, 17]. In the following we take advantage of the theoretical framework of pulsating fronts to examine the spatial dynamics of different pathogens spreading in a one-dimensional environment. We further assume that the host is constantly exposed to a single selective pressure (e.g., an antibiotic) but the type of selection (e.g., the identity of the drug) varies periodically in space. In a second step, we allow mutations between different pathogen genotypes and we analyse the evolution of a coalition of different pathogen genotypes. Finally, we examine the effect of demographic stochasticity on the speed of spreading epidemics when host population are assumed to be finite.

## 2 Model

We model the dynamics of a directly transmitted pathogen in a one-dimensional habitat. At time *t* and position *x*, the host population is divided into uninfected individuals, *S*(*t*, *x*), and infected individuals, *I*(*t*, *x*). We assume that dead hosts are immediately replaced by new susceptible hosts so that the total density of hosts is assumed to remain constant over space and time: *K* = *S*(*t*, *x*) + *I*(*t*, *x*). We focus on a scenario where the environment is divided into two different habitats where all the hosts are treated by either drug *A* or drug *B*. But the two types of hosts could also be due to the use of two different vaccines or, if we consider the spread of a phytopathogen in crop, by the use of two resistant varieties in different fields. We consider a simple spatial pattern where treatment varies periodically and we use *L* to denote the period of the spatial fluctuation of the treatment. Because all the hosts are treated by a drug we expect that the pathogens sensitive to both drugs will be rapidly outcompeted by resistant genotypes. We thus focus our analysis on the dynamics of three resistant pathogen genotypes circulating in the host population: (i) the density of hosts infected with the genotype only resistant to drug *A* is noted *I_a_*(*t*, *x*), (ii) the density of hosts infected with the genotype only resistant to drug *B* is noted *I_b_*(*t*, *x*) and (iii) the density of hosts infected with the genotype resistant to both drugs is noted *I_m_*(*t*, *x*) (*m* for multiresistance). Coinfection by different genotypes is not allowed and each genotype *i* is characterized by *β_i_*(*x*), the rate at which transmission occurs between infected and susceptible hosts after a contact at position *x*. The rate of transmission of the multiresistant genotype *β_m_* is independent of space because multiresistance implies that the rate of transmission is not affected by the treatment. In contrast, the rates of transmission *β_a_*(*x*) and *β_b_*(*x*) vary in space because we assume that treatment reduces transmission (without affecting the other life history traits). All the infections are assumed to end (because of clearance and/or increased mortality due to pathogen virulence) at a rate *α*. More precisely, we assume that *β_a_* (resp. *β_b_*) takes values *α* + *r* in locations treated by drug *A* only (resp. *B* only), and value *α* – *r* in locations treated by drug *B* only (resp. *A* only), see **Fig. 1**. This symmetry between the two specialists simplifies the following analysis of the model. Note, however, that we also examine a scenario when we introduce some asymmetry in the maximal growth rates of the two specialists in the **Supplementary Information**(section 1.2.1). Mutations may occur between these three genotypes and *μ_ij_* stands for the rate of mutation from genotype *i* to genotype *j*.

**Figure 1:**
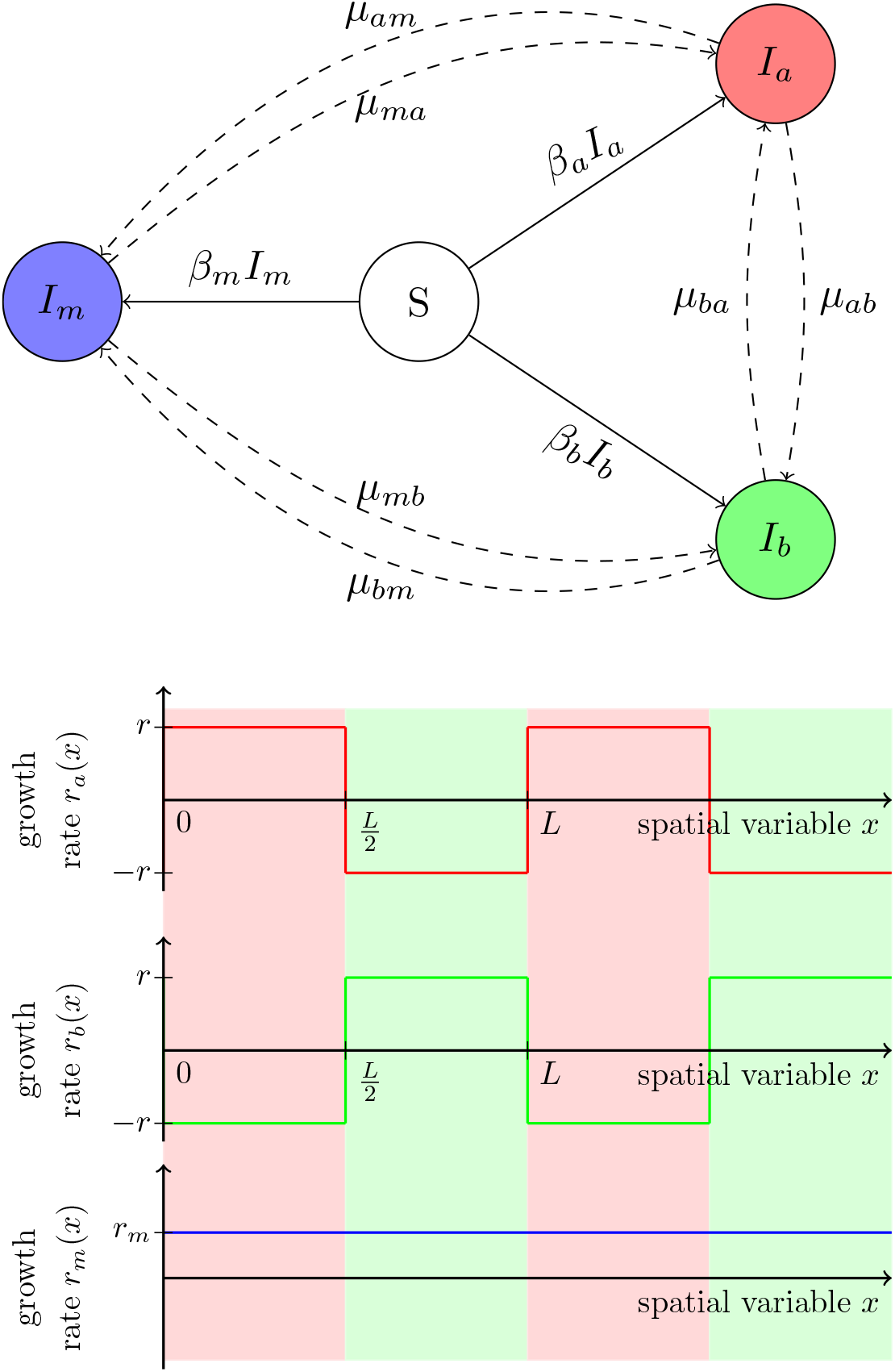
Schematic presentation of the evolutionary epidemiology model and the spatial heterogeneity of the environment. Top figure: diagram of the compartmental model. *S* represents susceptible hosts, *I_a_* (resp. *I_b_*, *I_m_*) represents hosts infected by the pathogen *a* resistant to drug *A* (resp. type *b* resistant to drug *B*, type *m*, resistant to both drugs *A* and *B*). In dashed we have represented mutations that typically happen at a much lower rate than transmissions. Bottom figure: Values of the intrinsic growth rates *x* ↦ *r_a_*(*x*) = *β_a_*(*x*) – *α*, *x* ↦ *r_b_*(*x*) = *β_b_*(*x*) – *α*, *x* ↦ *r_m_* = *β_m_* – *α* as a function of the spatial variable 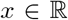, where, for *x* ∈ (0, *L*), 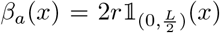 while 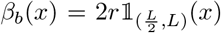, and *α* = *r*. The red (resp. green) area represents the locations 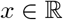 treated with drug *A* (resp. *B*).

The transmission of the pathogen is assumed to be local (infected hosts can only infect susceptible hosts at the same spatial location) but both susceptible and infected hosts are allowed to diffuse in one dimension with a fixed rate *σ*. In other words, we neglect the influence the pathogen may have on the mobility of its host. Our model can thus be written as the following set of reaction–diffusion equations (for readability, we drop the time and space dependence notation on host densities):

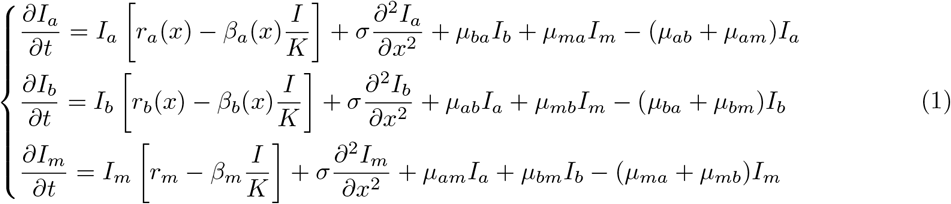

where *I* = *I_a_* + *I_b_* + *I_m_* and *r_i_*(*x*) = *β_i_*(*x*) – *α* is the malthusian growth rate of the single-resistance genotype *i* (with *i* ∈ {*a, b*}). Similarly, *r_m_* = *β_m_* – *α* is the malthusian growth rate of the multiresistant genotype *m*.

In the following we study the speed of spreading epidemics in a spatially heterogeneous environment as a function of (i) the period of the spatial fluctuation in drug use, (ii) the transmission rates of the different genotypes in the different habitats. We first consider the spread of single genotypes before analysing the effect of mutations among genotypes on the speed of a polymorphic pathogen population. Finally, we explore the effect of demographic stochasticity on the speed of monomorphic and polymorphic epidemics spreading in heterogeneous environments.

## 3 Results

### 3.1 The speed of a monomorphic pathogen population

The multiresistant genotype does not “feel” the spatial heterogeneity of the drug treatment. When such a genotype is introduced in the host population and if we assume no mutation (*μ_ma_* = *μ_mb_* = 0) the above system reduces to the spread of a single pathogen in a uniform environment. The pathogen population spreads as a traveling wave with a speed equal to [13, 14, 15]:

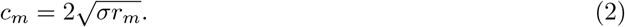

The analysis of the speed of a single-resistance genotype *i* ∈ {*a, b*} is more challenging because the growth rate of the pathogen varies periodically in space between *r_i_*(*x*) = *r* (when the genotype is resistant to the drug applied in *x*) and *r_i_*(*x*) = –*r* (when the genotype is not resistant to the drug applied in *x*). It is possible to derive good approximations for the speed of the epidemic in two limit cases [18, 19], namely when *L* is small and when *L* is large. When the period of the fluctuation of the environment is very small (i.e. *L* → 0) the *grain* of the environment is so small that the growth rate of the pathogen is equal to the average growth rate in the two habitats: 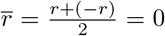. In contrast, when the period of the fluctuation is large the pathogen will move very fast when it is resistant to drug and it will slow down when the drug reduces its transmission rate. In the limit when *L* → ∞ the speed reaches an asymptote that can be described explicitly. We then get, for *i* ∈ {*a, b*},

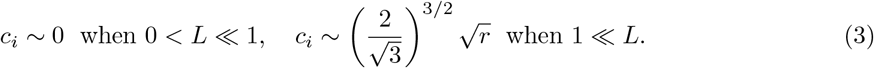

Moreover the speed of the single-resistance genotype epidemic increases with *L*, the period of the spatial fluctuation of the environment (**Fig. 2**).

**Figure 2:**
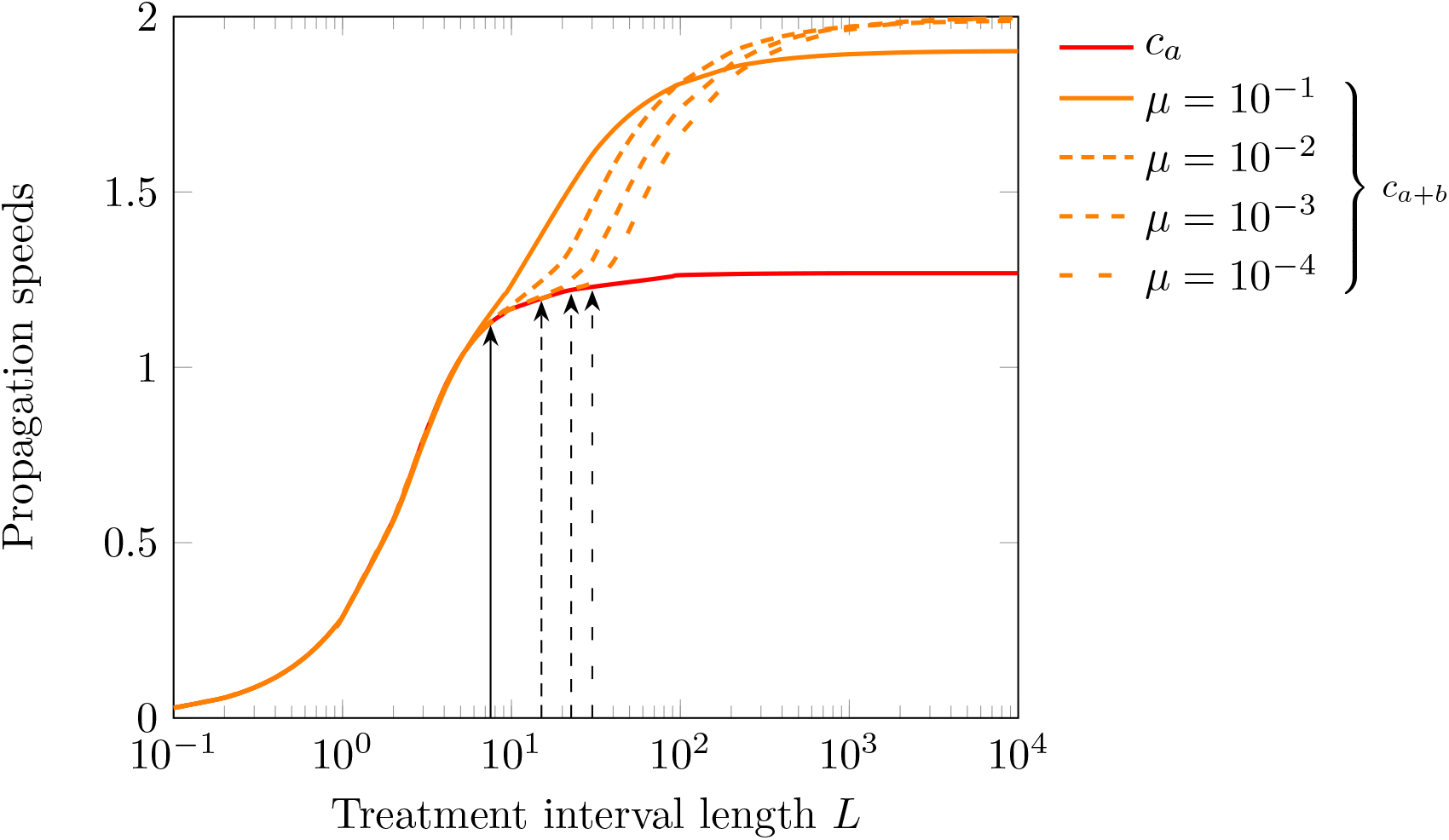
Propagation speed of single-resistance genotypes as a function of the period *L* and the mutation rate *μ*. We plot the speed *c_a_* of a single specialist genotype (red line) and the speed *c_a+b_* of a coalition of two specialists (orange lines) when *μ_ab_* = *μ_ba_* = *μ* (with *μ_am_* = *μ_bm_* = 0). The final values for *c_a_* are extrapolated (from *L* = 2000 inclusive). The black arrows indicate the values of *L_c_* for the different rates of mutation (see equation (4)). Parameters: *σ* = 1, *r* = 1, and the functions *β_a_*(*x*), *β_b_*(*x*), *r_a_*(*x*) and *r_b_*(*x*) are as in Figure 1.

### 3.2 The speed of a polymorphic pathogen population

Before considering the full system (with the 3 resistant genotypes) we examine the dynamics of a coalition of two single-resistance genotypes that resist two different drugs. When the mutation rates are very low (i.e. *μ_aj_* = *μ_bj_* ≈ 0) we recover the result of a monomorphic population (red line in **Fig. 2)**. However, numerical simulations with a fixed mutation rate *μ* between single-resistant genotypes indicate that increasing the mutation rate has a complex effect on the speed of the polymorphic population (**Fig. 2)**. When *L* is small, increasing the mutation rate has only a weak effect on epidemic speed because the environment changes so fast that both resistant genotypes are almost equifrequent. For intermediate values of *L*, the size of the area treated homogeneously with a single drug allows the resistant genotype to outcompete the other genotype and to take up some speed. Hence, the composition of the epidemic fluctuates between the two specialist genotypes and a higher mutation rate speeds up the emergence of this locally adapted genotype and increases the propagation speed. For larger values of *L*, however, this effect is dominated by the detrimental emergence of ill-adapted mutants (*mutation load*) that slows down the propagation within an area treated by a single drug. Hence, the composition of the pathogen population at the front of the epidemic depends on the balance between local selection, mutation and *L* which measures the amount of spatial heterogeneity. We show in the **Supplementary Information** (section 1.2.1) that there is a threshold value *L_c_* below which the whole epidemic can be driven by a single specialist:

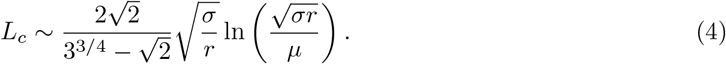

When *L* < *L_c_* the propagation of each specialist is independent because they can move through the “bad habitat” by diffusion. In contrast, when *L* > *L_c_* the bad habitat slows down the spread of the maladapted specialist and the coalition of two specialists is faster than a single specialist because they “pass the baton” when they move to a different habitat. The composition of the pathogen population at the front of the epidemic fluctuates between the two specialist genotypes. Higher mutation rates speed up the epidemic because mutation speeds up the switch between the two specialists at the tip of the front. Note, however, that high mutation rates generate a *mutation load* when *L* ≫ *L_c_* via the recurrent introduction of a drug sensitive genotype in the pathogen population. This is why the maximal speed of the coalition of single-resistance genotypes can never reach the speed of a universally adapted pathogen (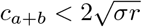 in **Fig. 2**).

When we assume a fixed mutation rate *μ* among the three resistant genotypes, the epidemic spreads faster than epidemics where only the coalition of two specialists is present, provided the period of the fluctuation is small (**Fig. 3**). Indeed, when *L* is small the multiresistant genotype outpaces the single-resistant genotypes at the front of the epidemic (**Fig. 3)**. In contrast, when *L* is large, the multiresistant genotype is outcompeted by the coalition of the two specialists (in particular when the mutation rate between single-resistant genotypes is large enough). Increasing the mutation rate tends to lower the speed of the epidemic when *L* is small or very large, because mutations reintroduce maladapted genotypes and build up the mutation load (**Fig. 4)**. For intermediate values of *L*, however, increasing the mutation rate can increase the speed of the pathogen spread, by speeding up the propagation of a the coalition of specialists *a* and *b* (**Fig. 4)**. This is due to the beneficial effects of mutations on the speed of the coalition of two single-resistant genotypes that we discussed above (**Fig. 2)**.

**Figure 3:**
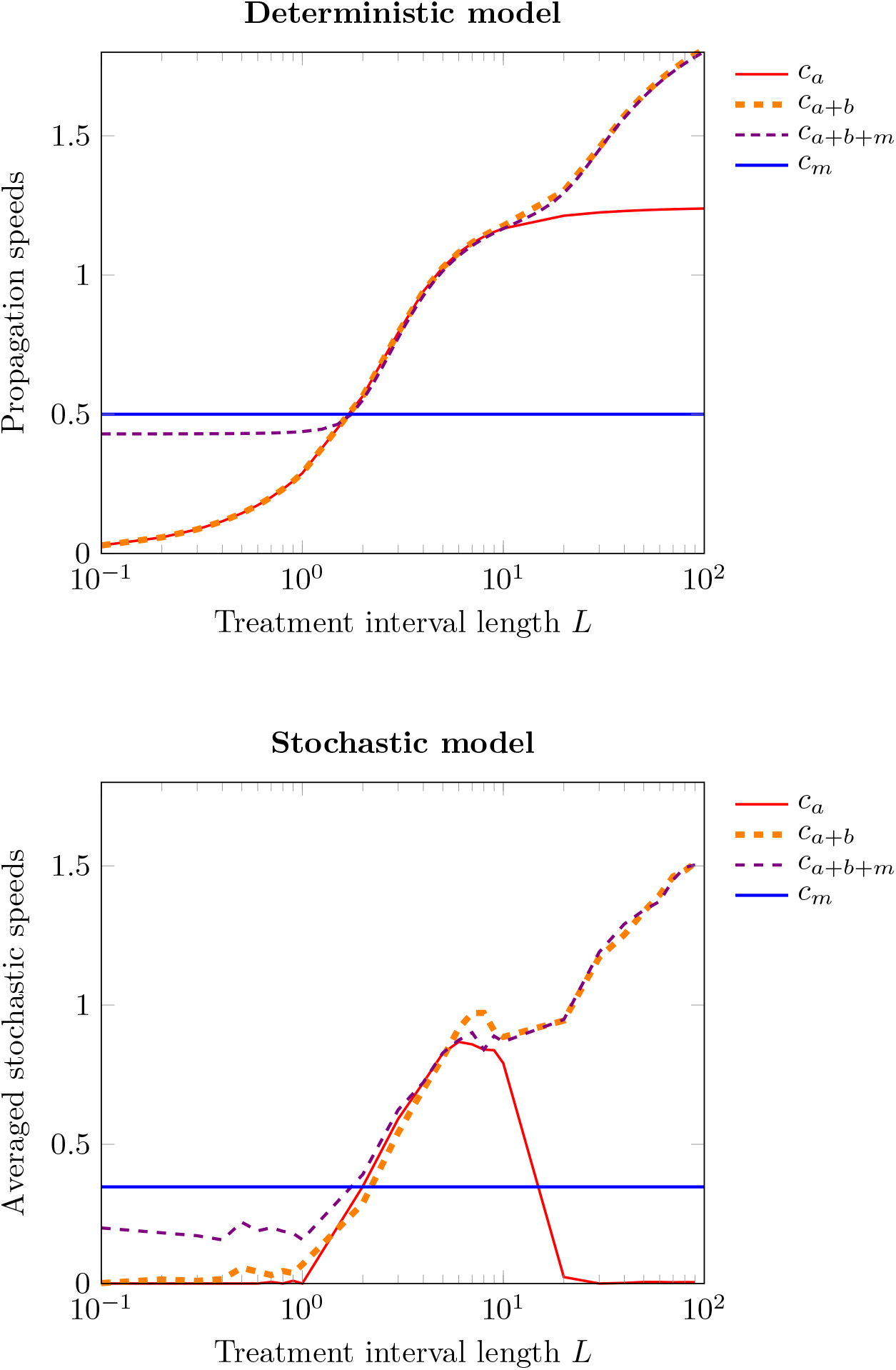
Propagation speed when only one specialist type is present (*c_a_*), both specialists are present (*c_a+b_* with *μ_ab_* = *μ_ba_* = *μ*) and when all the three resistant genotypes are present (*c_a+b+m_* with *μ_ij_* = *μ*, ∀ *i, j* ∈ {*a, b, m*}). Top figure: speed of the epidemic in the *deterministic model* (1) against the period *L* for the coalition of specialists (orange line: *c_a+b_* with *μ_ab_* = *μ_ba_* = *μ*), the generalist alone (blue line: *c_m_*) and the full model with both the specialists and the generalist (purple line: *c_a+b+m_* with *μ_ij_* = *μ*, ∀ *i,j* ∈ {*a, b, m*}). Bottom figure: speed of the epidemic in the *stochastic model* with *N* = 100 and *δx* = 0.1. Parameters: *r* = 1, 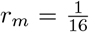, *σ* = 1, *μ* = 0.01, 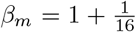, and the functions *β_a_*(*x*), *β_b_*(*x*), *r_a_*(*x*) and *r_b_*(*x*) are as in Figure 1.

**Figure 4:**
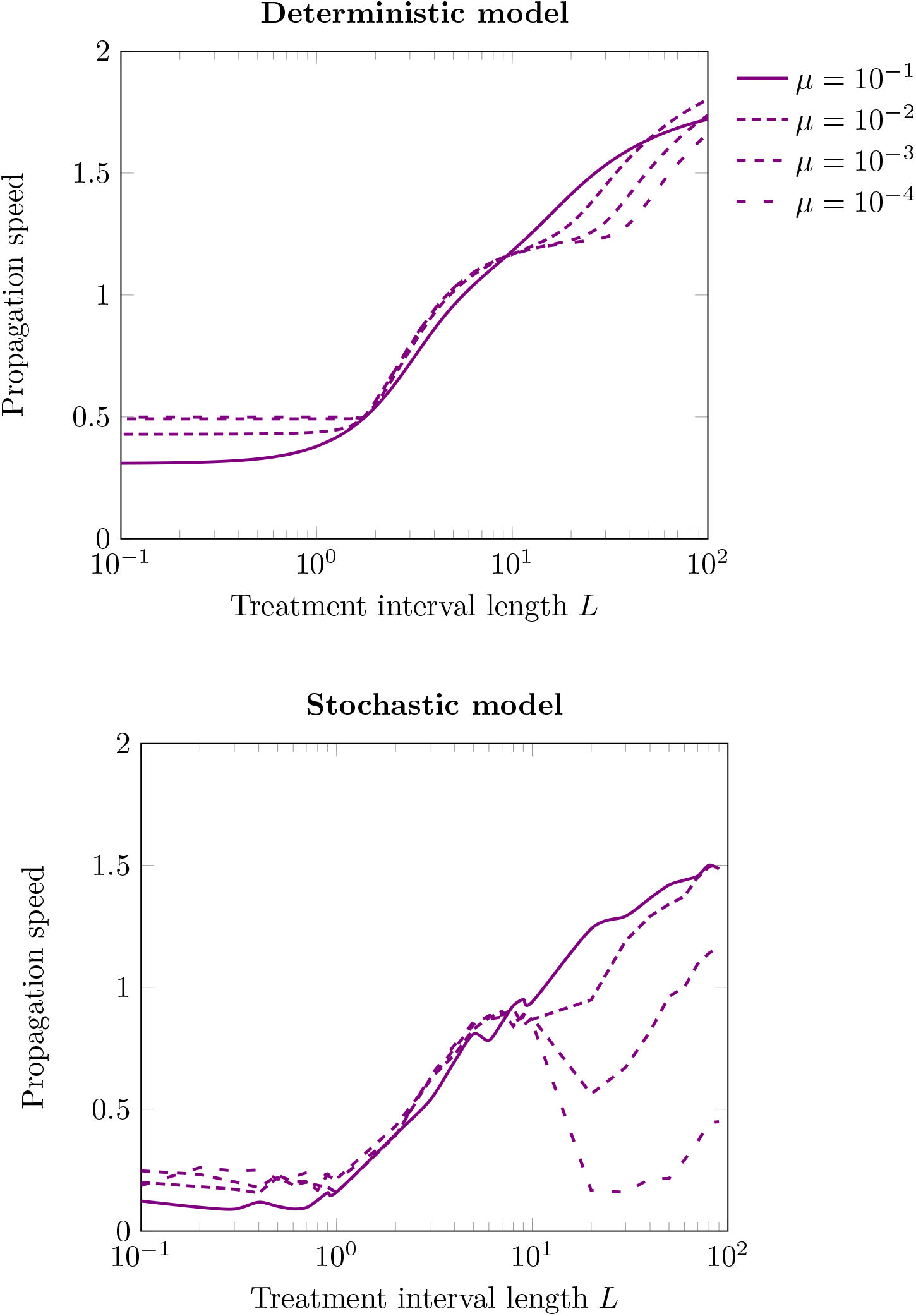
Effect of the mutation rate *μ* on the propagation speed of the epidemics when all three pathogen types are present (*c_a+b+m_* with *μ_ij_* = *μ*, ∀ *i, j* ∈ {*a, b, m*}). Top figure: *deterministic model*. Bottom figure: *stochastic model* with *N* = 100 and *δx* = 0.1. Parameters: *σ* = 1, 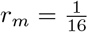, *r* = 1, 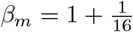, and the functions *β_a_*(*x*), *β_b_*(*x*), *r_a_*(*x*) and *r_b_*(*x*) are as in Figure 1.

### 3.3 The speed of stochastic epidemics

The above results rely on the assumption that the deterministic model we are using provides a good description of the spread of a pathogen epidemics. Yet, the front of the epidemic is necessarily driven by a small number of infections. The finite nature of the pathogen population at the edge of the epidemics yields demographic stochasticity and is expected to slow down its spread [14, 20, 21, 22]. In the following we explore the effect of stochasticity using an individual-based model that takes into account the finite number *N* of hosts at each spatial location. The individual transitions between the different states of the hosts are described by a list of random events (transmission, mutation, death; see the **Supplementary Information** for a detailed description of the individual-based model, section 2). As expected, this stochastic model converges to the above deterministic model when *N* is assumed to be very large. To study the effect of demographic stochasticity on epidemic spread we performed simulations with our individual-based model and measured the average speed on a long time interval after the influence of the initial condition is lost.

First, we discuss the speed of monomorphic epidemics in the absence of mutations. The speed of the multiresistant genotype is decreased by the effect of stochasticity but remains very close to the deterministic approximation (see [14, 20]). The magnitude of this drop is expected to be proportional to 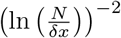, where 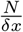 represents the number of hosts per unit of space. In contrast, the speed of the single-resistant mutant is dramatically altered by stochasticity (**Fig. 3)**. This speed is always lower than the speed of the deterministic approximation but, when *L* is large the speed can drop abruptly to zero which indicates that the pathogen cannot spread any more. Indeed, when the period of the fluctuation of the environment reaches a threshold value 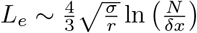 the pathogen cannot cross the unfavourable habitat (see **Supplementary Information**, section 2.2.2). In particular, the pathogen is very likely to go extinct in the unfavourable habitat when the population size is small, the diffusion rate is limited and its growth rate is very negative (remember that we assume the growth rate to be –*r* in the unfavourable habitat). Note that this critical period *L_e_* only increases logarithmically with the population size *N*, so that this blocking effect can be observed even with relatively large population sizes. This explains why the propagation speed of a single-resistance genotype is maximised for intermediate values of *L*. In the deterministic approximation, in contrast, the pathogen can always cross unfavourable habitats because extinctions do not occur and the speed of epidemic spread increases monotonically with *L*.

Second, if we allow some mutation between the two single-resistant genotypes, the epidemic can cross those unfavorable environments because mutations will rescue pathogen populations when *L* > *L_e_*. Consequently, increasing mutation rates can have a dramatic impact on the speed of epidemics when *L* is large (**Fig. 4**). Finally, when we allow the mutation between the three different genotypes, the speed of the epidemics is close to (but lower) than the deterministic approximation, and this speed can decrease when *L* > *L_e_* and the mutation rates are small enough (**Fig. 4)**. As pointed out above, the magnitude of this effect on the reduction of the epidemic speed is of the order (ln(*N*))^-2^ when *N* is large enough.

### 3.4 Pathogen diversity far behind the epidemic front

In the previous sections we focused on the composition of the pathogen population at the edge of the epidemic. Next, we characterise the composition of the pathogen population far behind the front, when it reaches an endemic equilibrium. Note that the composition of the pathogen population behind the front is much less sensitive to the effect of demographic stochastity because at the endemic equilibrium, the number of pathogen present is much larger than at the front of the epidemics, diminishing greatly the risk of genotype extinctions. Hence, we do not need to distinguish the deterministic and stochastic models in this section. Three cases can be observed (**Fig. 5)**:

i. **The multiresistant genotype dominates:** If both the cost of being multiresistant (i.e. *r* – *r_m_*) and *L* are low, the generalist strategy outcompetes the specialists and goes to fixation.
ii. **The coalition of specialist genotypes dominates:** When both the cost of being multiresistant (i.e. *r* – *r_m_*) and *L* are large, the coalition of specialists outcompetes the generalist strategy.
iii. **The three genotypes coexist:** The coexistence between the three different genotypes is also possible for a range of parameter values when both *r_m_* and *L* are relatively large. As pointed by [23], a generalist strategy can outcompete specialits at the interface between habitats.

**Figure 5:**
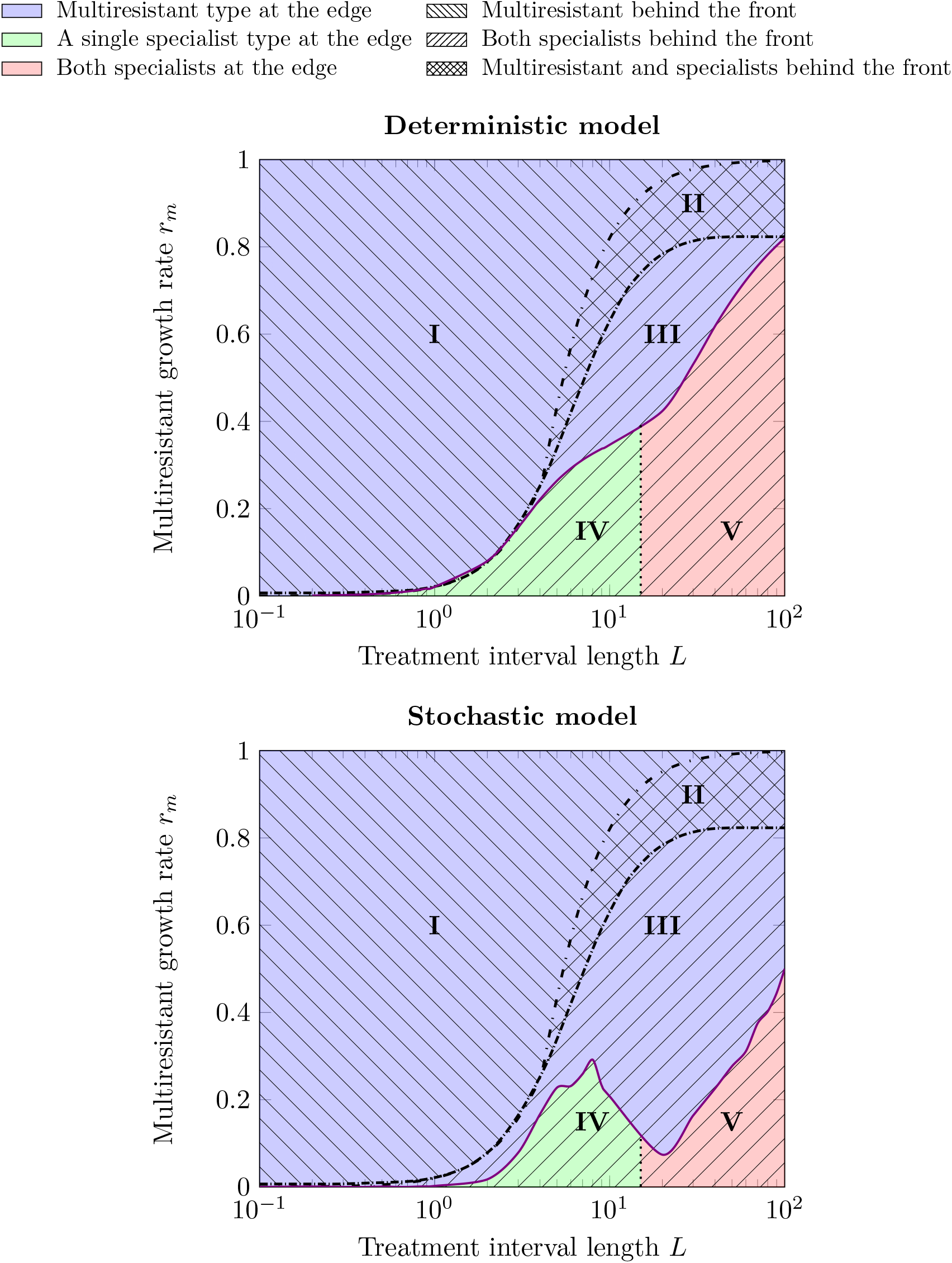
The five epidemic profiles. Composition of the population at the edge of the front (colors), and behind the front (hatches), as a function of *r_m_* and *L* with *μ_ij_* = *μ*, ∀ *i,j* ∈ {*a, b, m*}. See also **Fig. S2**, **S3** and **S6** in the **Supplementary Information** for the description of these different epidemic profiles. Top figure: *deterministic model*. Bottom figure: *stochastic model* with *N* = 100 and *δx* = 0.1. Parameters: *σ* = 1, *μ* = 0.01, *r* = 1, *β_m_* = 1 + *r_m_*, and the functions *β_a_*(*x*), *β_b_*(*x*), *r_a_*(*x*) and *r_b_*(*x*) are as in Figure 1.

### 3.5 Five epidemic profiles

Combining the different modes of propagation and the different types of pathogen populations behind the front, we can distinguish five different profiles of epidemics (**Fig. 5)**. Interestingly, we identify an epidemic type (marked by **III** in **Fig. 5**, see also **Fig. 6)** where the multiresistant genotype *m* drives the spread of the epidemic but is outcompeted later on by the coalition of the two specialists (single-resistance genotypes *a* and *b*). In other words, the analysis of the transitory dynamics reveals conditions where the multiresistant genotype is able to emerge, taking advantage of the presence of numerous uninfected host populations, even though specialized strategies are better competitors once the epidemics has developed and many hosts have been infected.

**Figure 6:**
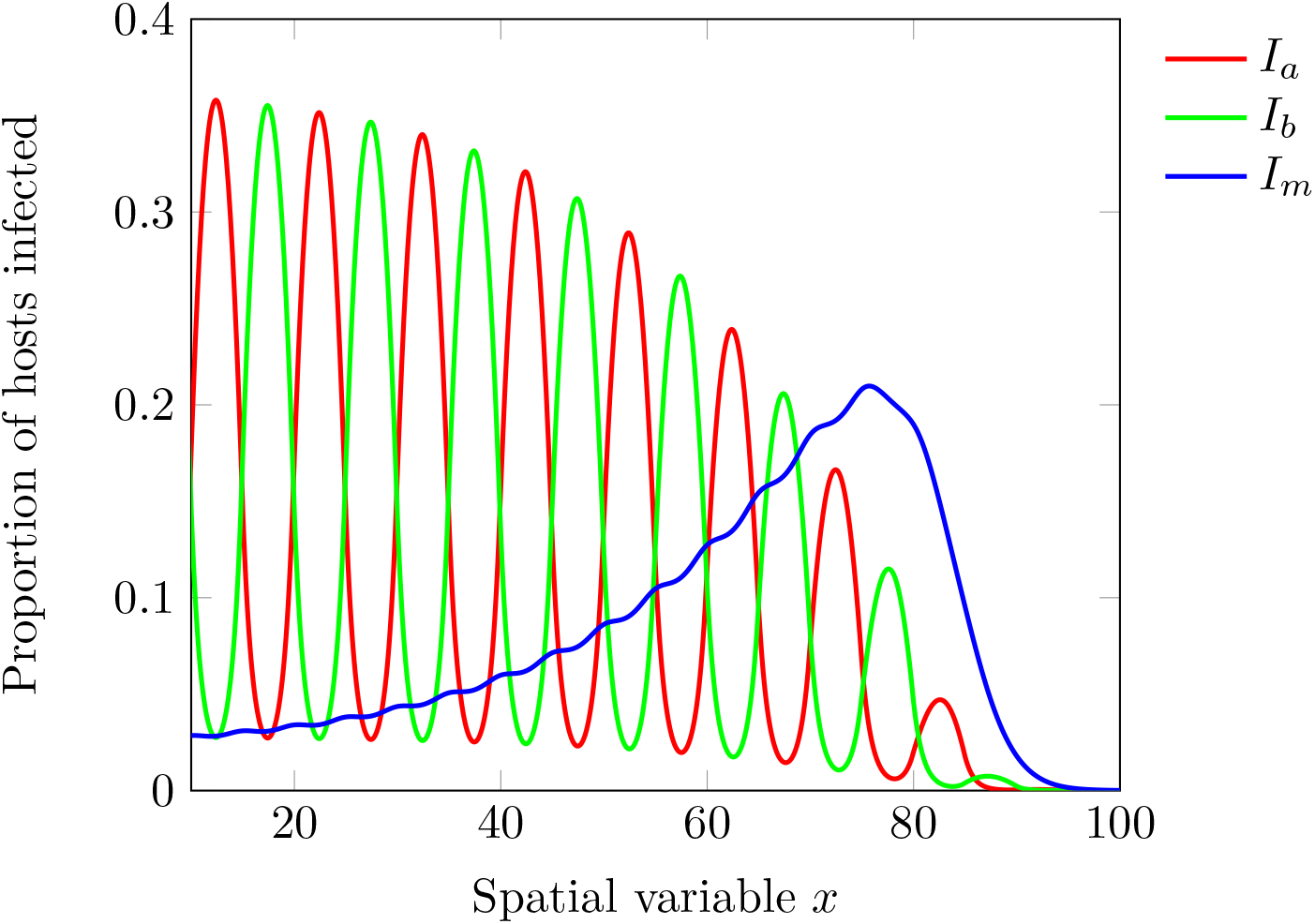
Composition of the pathogen population at the front of the epidemic. Propagating epidemics for *r_m_* = 0.5, *r* = 1, *μ* = 0.001, *L* = 10*β_m_* = 1.5, and the functions *β_a_*(*x*), *β_b_*(*x*), *r_a_*(*x*) and *r_b_*(*x*) are as in Figure 1. This corresponds to the epidemic type marked by **III** in **Fig. 5**.

We recover the same five epidemic profiles with finite host population sizes (**Fig. 5)** but demographic stochasticity affects the genetic diversity at the front of the epidemic where the size of the pathogen population is reduced. Single-resistance genotypes are most sensitive to the influence of stochasticity because these specialized genotypes can reach very low density in unfavourable habitats. The multiresistant genotype *m* benefits from the influence of this demographic stochasticity (compare the size of epidemic type marked by **III** in the deterministic and stochastic cases illustrated by **Fig. 5)**.

## 4 Discussion

Adaptation to a diverse range of selective pressures (e.g., drugs, specific immune responses, different host genotypes) is expected to result in the accumulation of multiple adaptations (drug resistance, immune escape mutations, virulence alleles) in pathogen populations. Ultimately, this process may lead to the emergence of multi-adapted (i.e. multiresistant) pathogens and a dramatic erosion of the efficacy of control measures. This major public-health concern requires a better understanding of the spatial spread of pathogen adaptations and our analysis challenges the view that the evolution of multiresistance is an unavoidable consequence of the use of multiple drugs against pathogens. We show that multiresistant pathogens can drive the spread of pathogen epidemics in spatially variable environments. Indeed, a multiresistant genotype is a *generalist* strategy that better copes with the heterogeneity of the environment during the early stage of a spreading epidemic. In the long term, however, the single-resistance genotypes are *specialist* strategies that are locally adapted to single drug treatments. Consequently, spreading epidemics may start with a multiresistant pathogen and end with a coalition of single-resistant genotypes (**Fig. 6)**.

Earlier studies have either looked at the evolution of polymorphic pathogen populations spreading in homogeneous environments [12, 13, 14] or at monomorphic populations spreading in heterogeneous environments [15, 16, 17]. The originality of our work relies on the analysis of the speed of a polymorphic population in a heterogeneous environment. We show that mutation is a double edged sword: (i) it allows the pathogen to acquire new resistance mutations and thus to adapt to new drugs but (ii) it can also produce a mutation load with the recurrent introduction of poorly adapted genotypes. The balance between these two effects depends on the heterogeneity of the environment which, in turn, depends on the ratio between the period *L* of the fluctuation of the environment and the diffusion coefficient *σ*. The beneficial effect of a higher mutation rate is maximal for intermediate levels of this ratio. Indeed, it is not profitable for the pathogen population to mutate often when the environment keeps changing (i.e., *L* ~ 0) or when the environment changes very slowly (i.e., *L* → ∞). Several earlier studies obtained similar conclusions in non-spatial models where it is possible to show that there is an optimal stochastic switching rate between specialized phenotypes that maximizes the growth rate of a population in a fluctuating environment [24, 25]. In all these different scenarios, the introduction of genetic variation provides a way to “pass the baton” between different specialist genotypes and allows the population to exploit more efficiently a fluctuating environment.

As expected from earlier theoretical studies [14, 20, 21, 22], demographic stochasticity lowers the speed of the epidemic spread. Most of the results of the deterministic model hold in finite host populations. The only notable exception occurs when large values of *L* can prevent the spread of single-resistance genotypes. The input of new mutations, however, may provide a way to adapt to the new drug. Hence the speed of pathogen epidemics may be constrained by both the stochastic nature of the demographic process and the stochastic nature of the mutation events occuring at the edge of the epidemic.

Several experimental studies have monitored and quantified the spread and the evolution of a bacteria in laboratory conditions [26, 27]. In particular, the MEGA-plate experiment of Baym et al followed the spread of *Escherichia coli* in a spatially heterogeneous environment characterised by increasing concentrations of antibiotics. This fascinating experiment allowed to visualize pathogen spread and evolution in real time. This experimental procedure could be used to test some of our predictions. For instance, we could monitor the influence of the scale of spatial heterogeneity with a manipulation of the parameter *L* in the MEGA-plate. We hope that the present theoretical framework may stimulate an experimental validation of our theoretical predictions using experimental evolution of microbes in spatially heterogeneous environments.

Our models can be used to make very practical recommendations regarding the manipulation of the spatial structure of the environment. This spatial structure can be manipulated by the spatially heterogeneous use of therapeutic drugs to treat infections [6, 28, 29] but also the deployment of resistance genes against pathogens in plants [7, 30, 31, 32, 33]. If the objective is to limit the speed of the epidemic spread, a lower value of *L* should be recommended. Lower *L* values imply that a spreading epidemics is exposed to a more variable environment. This prevents the pathogen to specialize to a specific environment and, consequently, to speed up in a favourable environment. Interestingly, fine-scale environmental heterogeneity (low *L* values) are also expected to reduce the probability of pathogen emergence [34]. This fine-scale heterogeneity, however, may promote the spread of generalist and multiresistant pathogens. Those pathogens are likely to spread more slowly because of the fitness cost associated with multiple resistance genes. Avoiding the spread of pandrug-resistance which erodes the efficacy of therapeutic drugs may be viewed as a public-health priority. Besides, after the spread of multiresistant genotypes, additional compensatory mutations (not considered in our model) may restore the competitivity of those genotypes against single-resistance genotypes. Those compensatory mutations could promote the persistence of pandrug-resistance in the long-term. In other words, the optimal deployment of control measures in space varies with the forecast horizon. Our model helps clarify the consequences of these interventions on the short term epidemiological dynamics (the speed of the spreading epidemic) as well as the evolutionary dynamics of the pathogen population.

## 5 Acknowledgements

We thank the CNRS MITI for funding the project VIRADAPT&SPREAD.

