## Supplementary Information for "Evolution and spread of multi-adapted pathogens in a spatially heterogeneous environment"

December 12, 2022

**A Remarks on the analysis method** **18**

**B Numerical estimates of the speed of the deterministic front** **18**

**C Probability of an individual crossing of an unfavourable zone** **20**

### 1 Deterministic model and the spread of drug resistance

#### 1.1 Model and existing propagation results

##### 1.1.1 The deterministic model

As described in the main text, at time $t$ and position $x$, the host population is divided into susceptible hosts, $S(t,x)$ and three genotypes of infected hosts: (i) hosts infected by a pathogen only resistant to drug A, $I_a(t,x)$, (ii) hosts infected by a pathogen only resistant to drug B, $I_b(t,x)$ and (iii) hosts infected by a pathogen resistant to both drugs, $I_m(t,x)$ ($m$ for multiresistant). We denote by $\beta_i(x)$ the rate at which transmission occurs between a host infected by the pathogen type $i$ to a susceptible host located in the same position $x$. All the infections are assumed to end (because of clearance and/or increased mortality due to pathogen virulence) at a rate $\alpha$. Mutations may occur between these three genotypes and $\mu_{ij}$ stands for the rate of mutation from genotype $i$ to genotype $j$. Finally, each host diffuses in space with a diffusion rate $\sigma$.

We are interested in an environment where the same amount of drugs A and B are applied in a periodic way, with a period $L$. At any point $x$ a single drug (A or B) is used and treatments are alternated in space in the following way: drug $A$ is present in location $x$ if $x \in \cup_{n\in\mathbb{Z}}[nL,(n+1/2)L)$, and drug $B$ is present in $x$ if $x \in \cup_{n\in\mathbb{Z}}[(n+1/2)L,(n+1)L)$. In other words, half of the environment is treated with drug A, while the remaining half is treated with drug B and $L$ measures the spatial heterogeneity of drug coverage. The impact of drug treatment appears in our setting through the transmission rates $x \mapsto \beta_a(x)$ and $x \mapsto \beta_b(x)$ that are then $L-$periodic. We assume that the multiresistant type is unaffected by either treatment A or B, so that $\beta_m(x) \equiv \beta_m$ is independent from location $x$.

If we consider a $1-$dimensional space and a large population, we obtain the following system of coupled reaction-diffusion equations:

$$\begin{cases} \frac{\partial S(t,x)}{\partial t} = \sigma\frac{\partial^2 S(t,x)}{\partial x^2} - S(t,x)\left(\frac{\beta_a(x)}{K}I_a(t,x) + \frac{\beta_b(x)}{K}I_b(t,x) + \frac{\beta_m}{K}I_m(t,x)\right) + \alpha I(t,x), \\ \frac{\partial I_a(t,x)}{\partial t} = \sigma\frac{\partial^2 I_a(t,x)}{\partial x^2} + \frac{\beta_a(x)}{K}S(t,x)I_a(t,x) - \alpha I_a(t,x) + \mu_{ma}I_m(t,x) + \mu_{ba}I_b(t,x) - (\mu_{ab}+\mu_{am})I_a(t,x), \\ \frac{\partial I_b(t,x)}{\partial t} = \sigma\frac{\partial^2 I_b(t,x)}{\partial x^2} + \frac{\beta_b(x)}{K}S(t,x)I_b(t,x) - \alpha I_b(t,x) + \mu_{mb}I_m(t,x) + \mu_{ab}I_a(t,x) - (\mu_{ba}+\mu_{bm})I_b(t,x), \\ \frac{\partial I_m(t,x)}{\partial t} = \sigma\frac{\partial^2 I_m(t,x)}{\partial x^2} + \frac{\beta_m}{K}S(t,x)I_m(t,x) - \alpha I_m(t,x) + \mu_{am}I_a(t,x) + \mu_{bm}I_b(t,x) - (\mu_{ma}+\mu_{mb})I_m(t,x). \end{cases} \tag{1}$$

For the sake of simplicity we assume that the total density of hosts is constant over space and time and equal to $K = S(t,x) + I_a(t,x) + I_b(t,x) + I_m(t,x)$. After dropping the time and space dependence notation on host densities, our model can be written in the following way:

$$\begin{cases} \frac{\partial I_a}{\partial t} = \sigma\frac{\partial^2 I_a}{\partial x^2} + I_a\big[r_a(x) - \frac{\beta_a(x)}{K}(I_a+I_b+I_m)\big] + \mu_{ma}I_m + \mu_{ba}I_b - (\mu_{ab}+\mu_{am})I_a \\ \frac{\partial I_b}{\partial t} = \sigma\frac{\partial^2 I_b}{\partial x^2} + I_b\big[r_b(x) - \frac{\beta_b(x)}{K}(I_a+I_b+I_m)\big] + \mu_{mb}I_m + \mu_{ab}I_a - (\mu_{ba}+\mu_{bm})I_b \\ \frac{\partial I_m}{\partial t} = \sigma\frac{\partial^2 I_m}{\partial x^2} + I_m\left[r_m - \frac{\beta_m}{K}(I_a+I_b+I_m)\right] + \mu_{am}I_a + \mu_{bm}I_b - (\mu_{ma}+\mu_{mb})I_m \end{cases} \tag{2}$$

with:

$$r_a(x) = \beta_a(x) - \alpha, \quad r_b(x) = \beta_b(x) - \alpha, \quad r_m = \beta_m - \alpha.$$

To simplify the notations and the analysis, we make further assumptions on parameters:

$$\beta_a(x) = \begin{cases} 2r & \text{if } x \in \cup_{n\in\mathbb{Z}}[nL,(n+1/2)L), \\ 0 & \text{if } x \in \cup_{n\in\mathbb{Z}}[(n+1/2)L,(n+1)L), \end{cases} \quad \beta_b(x) = \begin{cases} 0 & \text{if } x \in \cup_{n\in\mathbb{Z}}[nL,(n+1/2)L), \\ 2r & \text{if } x \in \cup_{n\in\mathbb{Z}}[(n+1/2)L,(n+1)L), \end{cases}$$

that is the pathogens resistant to treatment $A$ (and resistant to $A$ only) have a transmission rate equal to 0 if they are in an area treated with drug $B$, and a transmission rate $2r$ if they are in an area treated with drug $A$. If we assume moreover that the clearance/death rate of pathogens is independent of their position and equal to $r$, we obtain:

$$r_a(x) = \begin{cases} r & \text{if } x \in \cup_{n\in\mathbb{Z}}[nL,(n+1/2)L), \\ -r & \text{if } x \in \cup_{n\in\mathbb{Z}}[(n+1/2)L,(n+1)L), \end{cases} \quad r_b(x) = \begin{cases} -r & \text{if } x \in \cup_{n\in\mathbb{Z}}[nL,(n+1/2)L), \\ r & \text{if } x \in \cup_{n\in\mathbb{Z}}[(n+1/2)L,(n+1)L). \end{cases} \tag{3}$$

In other words, we assume that the two specialist pathogens have very symmetric phenotypes in the environments treated by drug A or B (but we illustrate the consequence of asymmetric phenotypes in Figure S3).

The transmission rate $\beta_m$ of the multiresistant pathogen is constant in space but is assumed to be lower that the maximal transmission rate $\beta_a(x) = 2r$ reached by a single resistant pathogen in a favourable environment.

The growth rate $r_m = \beta_m - \alpha$ then satisfies $r_m < r$, and $r - r_m = \max \beta_a - \beta_m$ can be seen as the *cost of multiresistance*. If the growth rate of the multiresistant is negative, the multiresistant type goes to extinction and the model reduces to a simpler model with 2 pathogen types only. We may therefore restrict our analysis to cases where $r_m \in (0, r)$.

##### 1.1.2 Propagation speed of a monomorphic population of multiresistant pathogens

We consider a situation where only the multiresistant type is present with no mutation (i.e. $\mu_{ij} = 0$ for $i, j \in \{a, b, m\}$). Since the multiresistant type is not affected by the drug treatments, the propagation of the multiresistant type alone corresponds to a propagation in a homogeneous environment, for which many results exist. The dynamics of a pathogen population initially present in a limited region ($S(0,\cdot) > 0$ on a compact interval and $S(0,\cdot) \equiv 0$ elsewhere) is well described by travelling front solutions. These are particular solutions where the pathogens propagate through space at a constant speed. We refer to [13] for a historical reference, and to [9] for travelling fronts with several pathogen types.

If we neglect the impact of mutations, the dynamics of the multiresistant type alone is described by the following equation:

$$\frac{\partial I_m}{\partial t} = \sigma \frac{\partial^2 I_m}{\partial x^2} + I_m \left[ r_m - \frac{\beta_m}{K} I_m \right]. \tag{4}$$

Ahead of the front, the dynamics of the solution of this non-linear model is guided by the following linearised equation:

$$\frac{\partial I_m}{\partial t} \approx \sigma \frac{\partial^2 I_m}{\partial x^2} + r_m I_m.$$

The particular solution $I_m(t,x) = \frac{1}{\sqrt{4\pi t}} e^{-\left(\frac{x^2}{4\sigma} - r_m t\right)}$ of this linear equation does provide an accurate description of the propagation speed that the model (4) exhibits: $I_m(t, 2\sqrt{\sigma r_m}) = \frac{1}{\sqrt{4\pi t}} \sim 1$. The propagation speed for a monomorphic multirestistant population, that is initially present in a bounded region only, is then:

$$c_m = 2\sqrt{\sigma r_m}. \tag{5}$$

If we take into account mutations in a homogeneous environment with different types, it was shown in [9] that the population propagates at a speed close to that of the fastest type. More precisely, the speed of the population is the speed of the fastest type reduced linearly by the mutation rates $\mu_{i,j}$ (for $i, j \in \{a, b, m\}$), when these are small.

##### 1.1.3 Propagation speed of a monomorphic population of single-resistant pathogens

We now consider a situation where the pathogen type resistant to drug A only is the only type present. Furthermore, we assume that there are no mutations (i.e. $\mu_{ij} = 0$ for $i, j \in \{a, b, m\}$), so that only the genotype $a$ is present at all times. By symmetry, equivalent results are obtained for genotype $b$. Since drug B is $L$-periodically present in the environment, the propagation of the type $a$ pathogen is akin to a propagation in a heterogeneous environment. In such an environment, we cannot expect the existence of proper travelling front solution because the heterogeneity of the environment breaks the self-similarity of travelling fronts. It is however possible to generalize the notion of travelling front for periodic media. The natural extension is the so-called *pulsating front*, first introduced by [17] in a biological context, and by Xin [21, 19, 20] in the framework of flame propagation. We also refer, among others, to the seminal works of Weinberger [18], Berestycki and Hamel [3]. Pulsating fronts for (2) were recently constructed in [1].

The equation satisfied by the type a pathogen is the following:

$$\frac{\partial I_a}{\partial t} = \sigma \frac{\partial^2 I_a}{\partial x^2} + I_a \left[ r_a(x) - \frac{\beta_a(x)}{K} I_a \right]. \tag{6}$$

Ahead of the front, this dynamics reduces to:

$$\frac{\partial I_a}{\partial t} \approx \sigma \frac{\partial^2 I_a}{\partial x^2} + r_a(x) I_a.$$

The speed of propagation of a localized initial datum for the above heterogeneous equation [17], [16], [5], [11], [12], satisfies a variational characterisation:

$$c_a = \min_{\lambda > 0} \frac{k_L(\lambda)}{\lambda}, \tag{7}$$

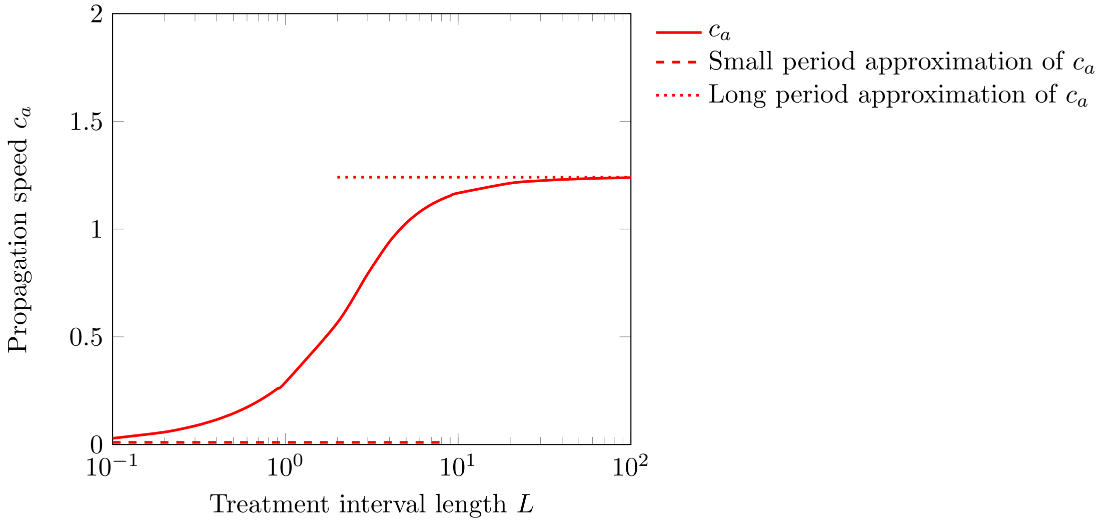

**Figure S1: Propagation speed of the deterministic model for a specialist alone.** Parameters: $r = 1$, $\sigma = 1$.

where $k_L(\lambda)$ denotes the periodic principal eigenvalue of an underlying second order operator (see (18)). This formula is not explicit, but a few approximations of this formula exist, when $L$ is small (for any periodic function $\beta_a$), and when $L$ is large (when $r_a(\cdot)$ is a square wave function). We detail these two approximations below.

**Small period approximation:** When the period of the fluctuation of the environment is very small (i.e. $L \to 0$) the *grain* of the environment is so small that the pathogen does not *feel* the spatial variation of the environment and its growth rate is the average growth rate in the two habitats: $\frac{1}{L}\int_0^L r_a(y)\,dy$. In this case, the speed of the single-resistance genotype is equal to:

$$c_a \xrightarrow[L\to 0]{} 2\sqrt{\frac{\sigma}{L}\int_0^L r_a(y)\,dy}.$$

In the particular case that we consider in this manuscript, $r_a$ is given by (3), and thus $c_a \to 0$ as $L \to 0$.

**Long period approximation:** In contrast, when the period of the fluctuation is large the pathogen population will grow within the favourable zones and the propagation will be limited by the difficult migration from one of these favourable patches to the next. In the limit when $L \to \infty$ the speed can be approximated for the type of growth function that we consider here ($r_a$ satisfying (3)): thanks to [11],

$$c_a \xrightarrow[L\to\infty]{} \sqrt{\sigma r}\left(\frac{2}{\sqrt{3}}\right)^{3/2}. \tag{8}$$

In Figure S1 we present a numerical computation of the propagation speed as a function of $L$. The approximations presented above are consistent with numerical results: $c_a \sim 0$ when $0 < L \ll 1$ and $c_a \sim \left(\frac{2}{\sqrt{3}}\right)^{3/2} \approx 1.24$ when $1 \ll L$ and $r = \sigma = 1$.

#### 1.2 Analysis of the deterministic model with some mutation

##### 1.2.1 Propagation speed for the coalition of single-resistance genotypes

In this section, we consider that the two specialists are present, but the multiresistant type is not. We assume that $\mu_{ab} = \mu_{ba} =: \mu > 0$, while $\mu_{am} = \mu_{bm} = 0$. Only the two specialist types are thus present in the system. We call this a coalition of single-resistance genotypes:

$$\begin{cases} \frac{\partial I_a}{\partial t} = \sigma\frac{\partial^2 I_a}{\partial x^2} + I_a\big[r_a(x) - \frac{\beta_a(x)}{K}(I_a + I_b)\big] + \mu I_b - \mu I_a, \\ \frac{\partial I_b}{\partial t} = \sigma\frac{\partial^2 I_b}{\partial x^2} + I_b\big[r_b(x) - \frac{\beta_b(x)}{K}(I_a + I_b)\big] + \mu I_a - \mu I_b. \end{cases} \tag{9}$$

We denote by $c_{a+b}$ the propagation speed of this system. Much less is known about the behaviour of the propagation speed with respect to $L$ in the case of polymorphic populations: we believe the propagation speed of (9) is equal to the propagation speed associated with the linearised system (see [1]), but even this property has not been rigorously proven yet. When linearised, (9) becomes

$$\begin{cases} \frac{\partial I_a}{\partial t} = \sigma \frac{\partial^2 I_a}{\partial x^2} + r_a(x) I_a + \mu I_b - \mu I_a, \\ \frac{\partial I_b}{\partial t} = \sigma \frac{\partial^2 I_b}{\partial x^2} + r_b(x) I_b + \mu I_a - \mu I_b. \end{cases} \tag{10}$$

From numerical computations (see Figure 2 in the main text), we see that the spread of the coalition may follow two types of propagation:

**When $L$ is small, $L < L_c$: *Independent propagation of the specialists***
A first way to propagate is to rely on the propagation of the specialists types independently: a specialist pathogen, e.g. pathogen $a$, thrives in the part of the environment treated by drug A and diffuses through the part of the environment treated by drug B to reach the next favourable area. Here are some properties of the *Independent propagation of the specialists*:

- Since the specialists propagate independently, mutations play a limited role and changing the mutation rate will have a limited effect on the propagation speed.
- If the symmetry between the $a$ and $b$ specialists breaks (e.g., if the maximal growth rate of the two specialists are slightly different, see Figure S3), the propagation would be driven by one specialist only.
- When it propagates, $I_a$ crosses areas treated by drug $B$ to reach the next favourable area. Most type $a$ individuals present in the areas treated with drug $B$ are the result of diffusion from favourable areas (rather than appearing from type $b$ parents through mutation). As a consequence, the value of $I_a$ in unfavourable areas is comparable to its level in neighbouring favourable areas.

The speed of the epidemics is close to the propagation speed of a specialist pathogen alone, that is $c_a$. We have then the following approximation when $\mu$ is small:

$$c_{a+b} \sim c_a. \tag{11}$$

More precisely, we expect that $c_{a+b}$ decreases linearly with $\mu > 0$ : mutations have a detrimental effect for this type of propagation.

**When $L$ is large, $L > L_c$: *Joint propagation of the specialists***
When $L$ is large, the propagation of the population through each interval of constant environment is driven by the corresponding specialist type. At the interface between intervals treated with different drugs, the composition switches from one type to the other thanks to mutations. Here are some properties of the *Joint propagation of the specialists*:

- Mutations are necessary at each interface between drug treatment to produce pathogens resistant to the drug present in the environment. The mutation rate has thus a big impact on the propagation speed: typically, a larger mutation rate leads to a faster propagation.
- If the symmetry between $a$ and $b$ is broken, typically if the maximal growth rate of the two specialists are slightly different (see Figure S3), the propagation will not be qualitatively impacted: the propagation will still rely on switches between the two specialist types.
- The pathogen population in the interval treated with drug A (resp. B) is dominated by the type $a$ (resp. $b$) specialists: the locally maladapted types have only a marginal presence if the mutation rates are small.

We propose next an explicit approximation of the speed for the *Joint propagation of the specialists*, when $L$ is large and $\mu$ is small. This argument is heuristic, and in future work it could be interesting to see if mathematically rigorous approximations could be produced. A natural way to address this problem would be to use Hamilton-Jacobi equations and Geometric Optics Approximations [8, 6].

We consider as an initial time $t_0$ the instant when the population of the type $a$ individuals reaches a size $\eta = \frac{\epsilon K L}{2}$ on the interval $[x_0, x_0 + L/2]$ treated with drug A (where $\epsilon < 1$ so that the density is far from the maximal density of the host and the linearised system is a good approximation for the front of the epidemics), that is

$$\int_{x_0}^{x_0+L/2} I_a(t_0, x)\, dx \sim \eta$$

We consider the situation where the infection is propagating from the left to the right of the line, and thus assume that at time $t_0$ the type $a$ pathogens are mainly concentrated on the left of the interval $[x_0, x_0 + L/2]$. We believe the distribution of individuals should relax towards the travelling wave profile of $I_a$ in a homogeneous favourable environment, which is approximately a decreasing exponential profile with parameter $\lambda = \sqrt{\frac{r}{\sigma}}$, that is

$$I_a(t_0, x) \sim \frac{\eta}{(1 - e^{-\sqrt{\frac{r}{\sigma}}\frac{L}{2}})}\sqrt{\frac{r}{\sigma}}e^{-\sqrt{\frac{r}{\sigma}}(x-x_0)}, \qquad \text{for } x \in [x_0, x_0 + L/2].$$

We can neglect the exponential term at the denominator right away since $L$ is large. Keeping this profile, the type $a$ individuals propagate through the favourable interval at a speed $2\sqrt{\sigma r}$ (we refer to Section 1.1.2 where we discuss the propagation of a pathogen in a strip of favourable environment). Then,

$$I_a(t, x) \sim \eta\sqrt{\frac{r}{\sigma}}e^{-\sqrt{\frac{r}{\sigma}}\left((x-x_0)-2\sqrt{\sigma r}(t-t_0)\right)},$$

before the competition has a large effect, that is for $x \in [x_0 + 2\sqrt{\sigma r}(t - t_0), x_0 + L/2]$. In particular, the number of type $a$ individuals present around $x_0 + L/2$ can be approximated by

$$I_a(t, x_0 + L/2) \sim \eta\sqrt{\frac{r}{\sigma}}e^{-\sqrt{\frac{r}{\sigma}}\left(L/2-2\sqrt{\sigma r}(t-t_0)\right)}.$$

These type $a$ individuals mutate at rate $\mu$ to type $b$ individuals, which are themselves able to grow at rate $r$ on the (almost empty) interval $[x_0 + L/2, x_0 + L]$. Then,

$$\frac{d}{dt}\int_{x_0+L/2}^{x_0+L} I_b(t, x)\,dx \sim \mu\eta\sqrt{\frac{r}{\sigma}}e^{-\sqrt{\frac{r}{\sigma}}\left(L/2-2\sqrt{\sigma r}(t-t_0)\right)} + r\int_{x_0+L/2}^{x_0+L} I_b(t, x)\,dx.$$

Solving this ordinary differential equation with the initial condition $\int_{x_0+L/2}^{x_0+L} I_b(t_0, x)dx = 0$ leads to

$$\int_{x_0+L/2}^{x_0+L} I_b(t, x)\,dx \sim \mu\eta e^{-\sqrt{\frac{r}{\sigma}}\frac{L}{2}}\frac{e^{2r(t-t_0)} - e^{r(t-t_0)}}{\sqrt{\sigma r}}.$$

Equipped with the above, we can compute the time $t_1$ when the number of individuals of type $b$ present on $[x_0 + L/2, x_0 + L]$ reaches $\eta$ and find

$$t_1 = t_0 + \frac{1}{r}\ln\left(\frac{1 + \sqrt{1 + 4\frac{\sqrt{\sigma r}}{\mu}e^{\sqrt{\frac{r}{\sigma}}\frac{L}{2}}}}{2}\right) \sim t_0 + \frac{1}{2r}\ln\left(\frac{\sqrt{\sigma r}}{\mu}e^{\sqrt{\frac{r}{\sigma}}\frac{L}{2}}\right) = t_0 + \frac{L}{4\sqrt{\sigma r}} + \frac{1}{2r}\ln\left(\frac{\sqrt{\sigma r}}{\mu}\right).$$

At time $t_1$, we can re-iterate the process above switching types $a$ and $b$. This propagation strategy then leads to the propagation speed

$$c_{a+b} \sim \frac{L/2}{t_1 - t_0} \sim \frac{2\sqrt{\sigma r}}{1 + \frac{2}{L}\sqrt{\frac{\sigma}{r}}\ln\left(\frac{\sqrt{\sigma r}}{\mu}\right)}$$

that is

$$c_{a+b} \sim 2\sqrt{\sigma r} - 4\frac{\sigma}{L}\ln\left(\frac{\sqrt{\sigma r}}{\mu}\right). \tag{12}$$

**Approximation of the critical length $L_c$**

We have described above two propagation strategies. Both are attempted by the propagating epidemics, and the fastest of these strategies will drive the front. Formula (11), associated to (8) (we assume here that $\mu > 0$ is small) describes the propagation speed of the first strategy, whereas the speed of the second strategy is approximated by (12). We can compare the propagation speeds of these two strategies and show that the first strategy is prevalent for $L < L_c$, while the second is prevalent for $L > L_c$. The transition between these two propagation strategies will correspond to the critical length $L = L_c$ such that the propagation speeds given by (11) and (12) are equal. Hence $L_c$ satisfies

$$\sqrt{\sigma r}\left(\frac{2}{\sqrt{3}}\right)^{3/2} = \frac{2\sqrt{\sigma r}}{1 + \frac{2}{L_c}\sqrt{\frac{\sigma}{r}}\ln\left(\frac{\sqrt{\sigma r}}{\mu}\right)},$$

that is

$$L_c = \frac{2\sqrt{2}}{3^{3/4} - \sqrt{2}}\sqrt{\frac{\sigma}{r}}\ln\left(\frac{\sqrt{\sigma r}}{\mu}\right). \tag{13}$$

##### 1.2.2 Propagation speed for the full deterministic model

In this section, we consider the full system (2), where both specialists and the multiresistant types are present. We assume that $\mu_{ij} = \mu > 0$ for $i, j \in \{a, b, m\}$. Numerical simulations (see Figure 3 in the main text) indicate that the propagation speed of solutions of the full system can be qualitatively approached by the highest propagation speed among the speed $c_m$ (propagation speed of the mutant alone) and $c_{a+b}$ (propagation speed of the coalition of specialists, when the mutation rate $\mu > 0$ is very small. We believe this idea provides a valuable qualitative description of the dynamics even if $\mu > 0$ is of the order of $\mu \in [10^{-4}, 10^{-1}]$ as considered in our simulations (see Figure S4). We believe it would be interesting to give a mathematical formulation of this idea in future work, to obtain a better understanding of this approximation and its limitations.

The effects of the growth rate of the multiresistant pathogen $r_m$ and the heterogeneity of the environment parameter $L$ on the composition of the pathogen population at the edge of the epidemic front is summarized in Figure S2:

- The front is driven by the multiresistant type if the propagation speed of the multiresistant type is greater than the propagation speed of the coalition, i.e. if $c_m > c_{a+b}$. This corresponds to the blue area in Figure S2, and to $L \in (0, 4.3)$ in Figure 3.
- The front is driven by the *independent propagation of the specialists* if $c_m < c_{a+b}$ and if the propagation speed of a specialist type alone is similar to the propagation speed of the coalition of specialists, i.e. $c_a \sim c_{a+b}$. This corresponds to the green area in Figure S2, and to $L \in (4.3, 10)$ in Figure 3.
- The front is driven by the *joint propagation of specialists* if $c_m < c_{a+b}$ and if the propagation speed of a specialist type alone is smaller than the propagation speed of the coalition of specialists, i.e. $c_a < c_{a+b}$. This corresponds to the pink area in Figure S2, and to $L \geq 10$ in Figure 3.

We illustrate a propagation driven by the multiresistant type in Figure S2(a). As described in Section 1.2.1, it is more difficult to observe the difference between the two other types of propagation. The difference would appear if the symmetry between the $a$ and $b$ types were broken. We represent in Figure S2(b) a situation where the front is driven by the *independent propagation of the specialists*, and in Figure S2(c) a situation where the front is driven by the *joint propagation of specialists.*

A higher mutation rate leads to the creation of ill-adapted pathogen types. Figure S4 shows that this is true in particular at the edge of an advancing epidemics. The effect of an increase in mutation rate on the speed of the epidemic front is a bit more complex: when the propagation is driven by a single pathogen type (the multiresistant type) or by the *Independent propagation of specialists*, increasing the mutation rates slows down the propagation speed because mutations generate mutants that are not helpful for the propagation. This corresponds to $L \leq 10$ in Figure S5. The same slowing down effect is present for $L$ very large, when the time necessary to propagate in favourable strips is the dominant factor determining the propagation speed of a coalition of specialists. This corresponds to $L \gtrsim 60$ in Figure S5 for $\mu = 10^{-1}$ and $\mu = 10^{-2}$. It would be interesting to see this effect for larger $L$ and smaller $\mu$, but the computational cost of such small mutation rates and large $L$ becomes prohibitive. We refer to Figure 2, where we have considered $L \in [10^{-1}, 10^{4}]$ and different mutation rates $\mu > 0$: we see in that figure that the propagation rate is unfavourably affected by $\mu > 0$ when $L$ is very large. When the coalition of specialists is driving the propagation, mutations enable the coalition to adapt to the alternating drug treatments (see Section 1.1.3 when $L > L_c$), and if $L$ is not very large, these mutations are actually the limiting factor deciding the propagation speed of the front. A higher mutation rate speeds up the propagation, which we can observe in Figure S5 for $L \in [10, 60]$.

##### 1.2.3 Composition of the population behind the front

In the previous section we focused on the analysis of the edge of the epidemic: we have analysed the propagation speed of the epidemics and, as a consequence, we have been able to discuss the composition of the pathogen population at the edge of the epidemics. We are now considering the composition of the pathogen population once the pathogen are settled, behind the epidemic front.

Numerical simulations show that the distribution of the pathogen population stabilizes into a single steady-state for the deterministic model (2). As illustrated by Figure S6, there are three families of possible steady-states:

- Steady states where only the generalist is present in large numbers. See Figure S6(a), where the multiresistant genotype outcompetes the specialist genotypes.
- Steady-states where all three genotypes coexist, as can be seen in Figure S6(b).

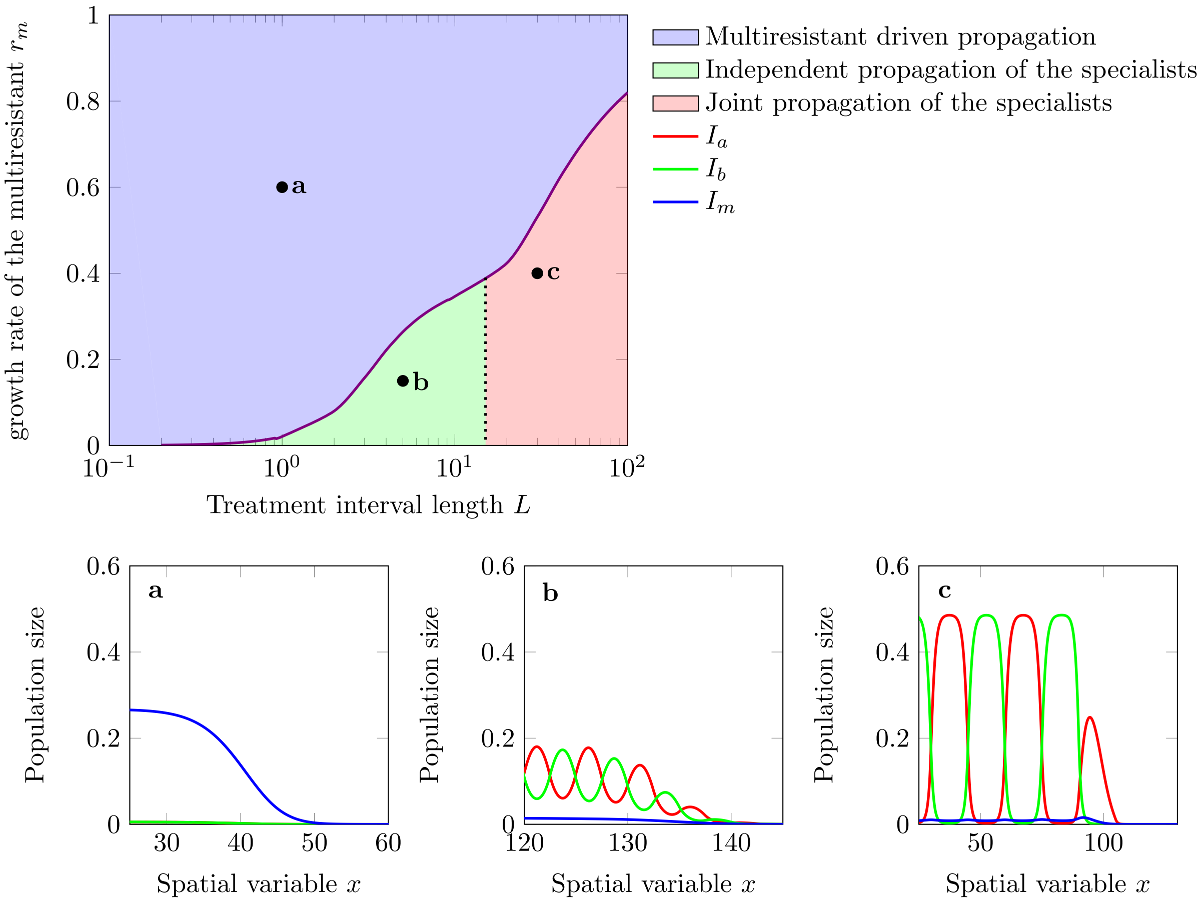

**Figure S2: Composition of the population at the edge of the epidemics.** The black line represents the coefficients $(L, r_m)$ for which the speed of the coalition of specialists is equal to the one of the multiresistant. The dark dotted line represents the coefficients $(L, r_m)$ where the speed of the coalition of specialists is equal to the propagation speed of a specialist type alone. The blue area (panel (a)) corresponds to a propagation led by multiresistant type. The green area (panel (b)) corresponds to a propagation led by the *independent propagation of the specialists* (i.e., mutations between the two specialists do not speed up the epidemic spread, see also equation (11)). The pink area (panel (c)) corresponds to a propagation led by the *joint propagation of the specialists* (i.e., mutations between the two specialists speed up the epidemic spread, see also equation (12)) . The effect of the transition between the green and the pink area on the composition of the front of the epidemic is more apparent when the maximal growth rates of the specialists are slightly different as illustrated in Figure S3 . Parameters: $\mu = 0.01$, $r = 1$.

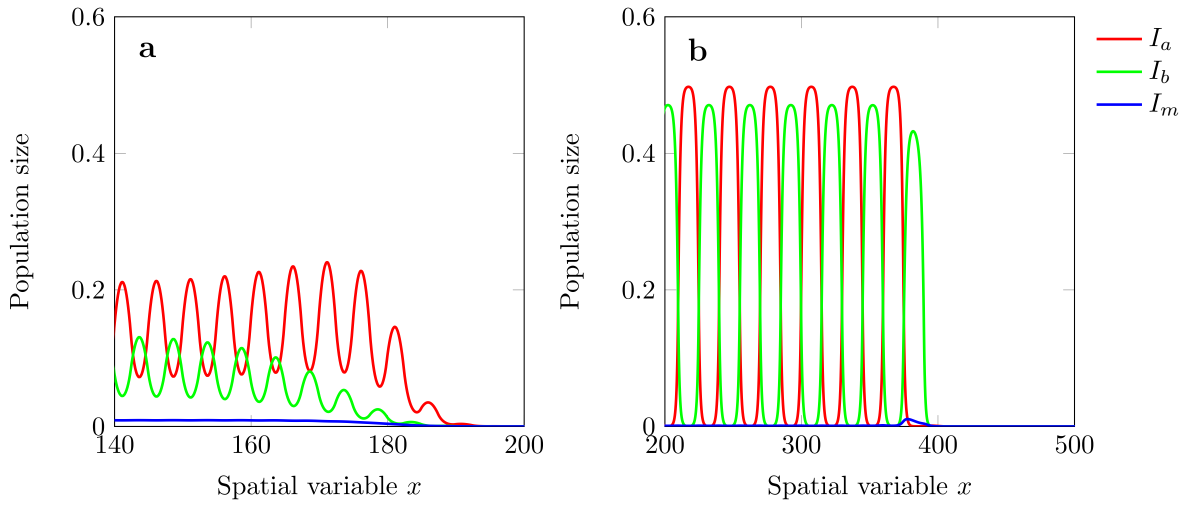

**Figure S3: Composition of the population at the edge of the epidemics when the two specialists have asymmetric growth rates.** The above graphs were obtained with the same coefficients as Figure S2(b) and Figure S2(c), except for $\beta_b$ reduced from 2 to 1.9 in favourable areas (intervals treated with drug $B$). This modification breaks the symmetry between the two specialists and has a large effect on the propagation dynamics. In (a) the propagation is led by $I_a$ only, and the other specialist (i.e,. $I_b$) only appears later on through mutation once the epidemics has settled. This is typical when the epidemics is led by the *independent propagation of the specialists* (see Figure S2). In (b), on the contrary, we see that both specialists are present at the front of the epidemic. In this case, the speed of the polymorphic population benefits from the mutation between specialists (see equation (12)) even if one of the two has a slightly slower growth rate. This is typical when the epidemics is lead by the *joint propagation of the specialists* (see Figure S2). Note that the asymmetry in growth rate also affects the composition of the pathogen population far behind the front (the $I_a$ strains reaches higher density than the $I_b$ strain) but both strains coexist.

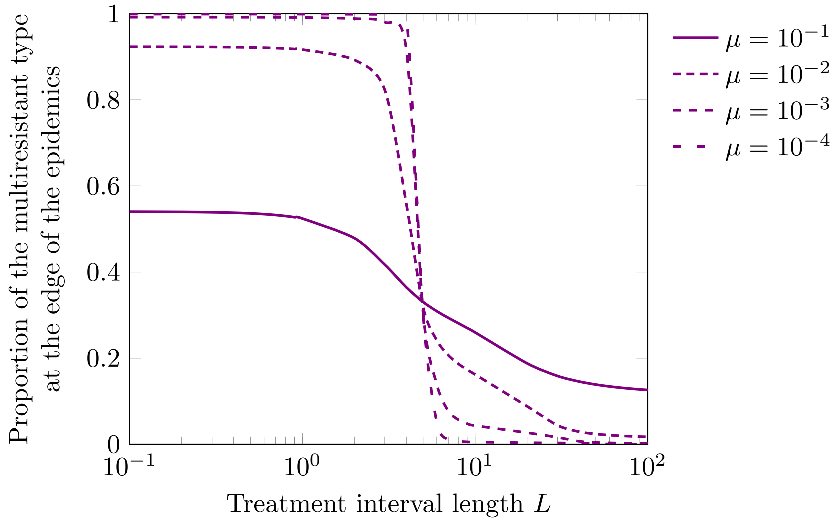

**Figure S4: Effect of the mutation rate $\mu$ on the proportion of the multiresistant type at the edge of the epidemics for the full deterministic model.** Parameters: $\sigma = 1$, $r_m = \frac{1}{4}$, $r = 1$.

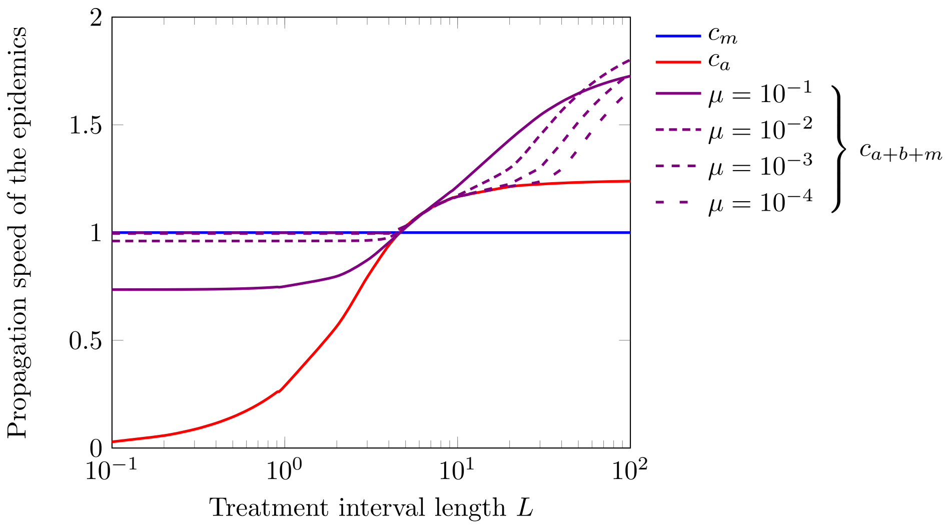

**Figure S5: Effect of the mutation rate $\mu$ on the propagation speed of the full deterministic model.** Parameters: $\sigma = 1$, $r_m = \frac{1}{4}$, $r = 1$.

- Steady-states where the two specialists are present in large numbers but the multiresistant type is only residual. This situation is illustrated by Figure S6(c).

In the following, we show how we can understand which of these steady-states is selected for given parameters.

The stability of a steady-state where only the multiresistant type is present is directly related to the analysis of the ability of a specialist genotype to invade that steady-state. We consider then a situation where only the multiresistant strain is present in the environment: $I_m(t,x) = \frac{r_m}{\beta_m} K$, while $I_a \equiv I_b \equiv 0$ (we neglect here the effect of mutations). If small populations of specialists $a$ and $b$ are introduced, their dynamics is given by the following linearised parabolic equations:

$$\begin{cases} \frac{\partial I_a}{\partial t}(t,x) - \sigma \frac{\partial^2 I_a}{\partial x^2} \sim I_a \left[ r_a(x) - \frac{\beta_a(x)}{K} I_m \right] = I_a \left[ -r + \mathbf{1}_{\cup_n \in \mathbb{Z}[nL,(n+1/2)L)} \left( 2r - \frac{2r\, r_m}{\beta_m} \right) \right], \\ \frac{\partial I_b}{\partial t}(t,x) - \sigma \frac{\partial^2 I_b}{\partial x^2} \sim I_b \left[ r_b(x) - \frac{\beta_b(x)}{K} I_m \right] = I_b \left[ -r + \mathbf{1}_{\cup_n \in \mathbb{Z}[(n+1/2)L,(n+1)L)} \left( 2r - \frac{2r\, r_m}{\beta_m} \right) \right]. \end{cases}$$

The growth of the specialist population $I_a$ is then related to the sign of the principal eigenvalue of the operator $\sigma \frac{\partial^2 I_a}{\partial x^2} + I_a \left[ r_a(x) - \beta_a(x) \frac{r_m}{\beta_m} \right]$. If we consider fixed parameters $r$ and $L$, we can use the monotony of the principal eigenvalue with respect to the growth rate to show that the principal eigenvalue associated to the operator above is positive as soon as $r_m \geq r_m^I$, where $r_m^I$ is such that there exist a positive solution $I_a$ to

$$\begin{cases} 0 = \sigma \frac{\partial^2 I_a}{\partial x^2}(x) + I_a(x) \left[ r_a(x) - \beta_a(x) \frac{r_m^I}{\beta_m} \right], \quad x \in \mathbb{R}, \\ I_a : \mathbb{R} \to \mathbb{R}_+, \quad L - \text{periodic}. \end{cases}$$

We can use formula (4.4) in [16] to provide a more analytic formula for $r_m^I > 0$: it is the smallest positive solution of the equation

$$\sqrt{1 - \frac{2r_m^I}{r + \bar{r}_m}} \tan \left( \frac{\sqrt{1 - \frac{2r_m^I}{r + r_m^I}} L}{2\sqrt{\sigma}} \right) = \tanh \left( \frac{\sqrt{r} L}{2\sqrt{\sigma}} \right).$$

We have shown that the steady-state composed of the mutliresistant type only is unstable if and only if $r_m < r_m^I$. It indicates that the specialist types are then present in the stable steady-state of the full model if and only if $r_m < r_m^I$. We represent this by north-east hatches in Figure S6. The steady-state composed of the mutliresistant type only is stable if $r_m^I < r_m$.

The stability of a steady-state composed of both specialist types only corresponds to the ability of a generalist pathogen to invade an environment where only the two specialist types are present. We denote by $(\bar{I}_a, \bar{I}_b)$ the steady distributions of the two specialists $a$ and $b$ when the multiresistant is absent (and if we neglect mutations), that is:

$$\begin{cases} 0 = \sigma \frac{\partial^2 \bar{I}_a}{\partial x^2}(x) + \bar{I}_a(x)\big[r_a(x) - \frac{\beta_a(x)}{K}\left(\bar{I}_a(x) + \bar{I}_b(x)\right)\big], \\ 0 = \sigma \frac{\partial^2 \bar{I}_b}{\partial x^2}(x) + \bar{I}_b(x)\big[r_b(x) - \frac{\beta_b(x)}{K}\left(\bar{I}_a(x) + \bar{I}_b(x)\right)\big], \\ \bar{I}_a,\, \bar{I}_b : \mathbb{R} \to \mathbb{R}_+, \quad L-\text{periodic}. \end{cases}$$

If a small population of multiresistants is introduced in this environment, its dynamics is governed by the following linearised parabolic equation:

$$\frac{\partial I_m}{\partial t}(t,x) - \sigma \frac{\partial^2 I_m}{\partial x^2}(t,x) \sim I_m(t,x)\left[r_m - \frac{r + r_m}{K}\left(\bar{I}_a(x) + \bar{I}_b(x)\right)\right],$$

and the population of multiresistant will survive if an only if the principal eigenvalue associated to this equation is positive. The maximum principle can be used to show $\bar{I}_a(x)+\bar{I}_b(x) \leq K$, and $r_m \mapsto r_m - \frac{r+r_m}{K}\left(\bar{I}_a(x) + \bar{I}_b(x)\right)$ is thus non decreasing. For fixed parameters $r$ and $L$, the monotony of the principal eigenvalue can then be used to show that the steady-state composed of the specialist types only is stable if and only if $r_m \leq r_m^{II}$, where $r_m^{II}$ is such that there exists a positive solution to the following elliptic eigenvalue problem:

$$\begin{cases} 0 = \sigma \frac{\partial^2 I_m}{\partial x^2}(x) + I_m(x)\left[r_m^{II} - \frac{r_m^{II}+r}{K}\left(\bar{I}_a(x) + \bar{I}_b(x)\right)\right], \quad x \in \mathbb{R}, \\ I_m : \mathbb{R} \to \mathbb{R}_+, \quad L-\text{periodic}. \end{cases}$$

We have shown that the steady-state composed of the specialist types only is unstable if and only if $r_m^{II} < r_m$. It indicates that the multiresistant type is present in the stable steady-state of the full model if and only if $r_m^{II} < r_m$. We represent this by north-west hatches in Figure S6. The steady-state composed of the specialists types only is stable if $r_m < r_m^{II}$.

For fixed parameters $r$ and $L$, if $r_m > r_m^I$, only the multiresistant is present in the steady-state of the full system, when the mutation rates are very small. If $r_m < r_m^{II}$, the two specialists are the only types present in the steady-state of the full system when mutation rates are very small. This corresponds in Figure S6 to the area with both types of hatches. Note mutations continuously produce residual populations of all types, but classification above remain a good qualitative description of the populations behind the fronts, when mutation rates are not too large.

### 2 Stochastic model of the spread of multiple drug resistance

In the above section, the derivation of the model (2) relies on the assumption that an infinite number of individual hosts are present at each spatial location. We relax this assumption in the following section and we explore the effect of a finite host population size on the speed and the composition of the pathogen population.

#### 2.1 The Individual Based Model

We consider a population of hosts that is distributed on a lattice $\delta x\,\mathbb{Z}$ for some $\delta x > 0$ (Figure S7). At each node of this lattice, there is a finite and constant number $N$ of hosts . With rate $\frac{\sigma}{2N\delta x^2}$, each couple of neighboring hosts (in the sense that one host is in $i\,\delta x$ and the other in $(i+1)\,\delta x$ for some $i \in \mathbb{Z}$) switch position. Each individual host may be infected by one of three types of infections: two *specialist* pathogens (denoted by $a$ and $b$) and a *multiresistant* pathogen (denoted by $m$). The infections are attached to their host: a host that switches to a different location carries its pathogens along. Moreover, each infected individual can recover or die at rate $\alpha$ (if the host dies it is immediately replaced by a new susceptible host so that the number of hosts remains constant to $N$). Transmission events occur within a lattice node (transmission is local in space). A host located in $i\delta x$ infected by a type $a$ pathogen (resp.$b$, $m$) initiates a transmission at rate $\beta_a(i\delta x)$ (resp. $\beta_b(i\delta x)$, $\beta_m(i\delta x)$). The receiver of this transmission is selected uniformly within the individuals present at the same location. The transmission occurs if the receiver is not carrying any pathogen prior to this transmission attempt (we assume that superinfections are not possible). Finally, each infection type mutates to each of the two other pathogen types with rate $\mu$.

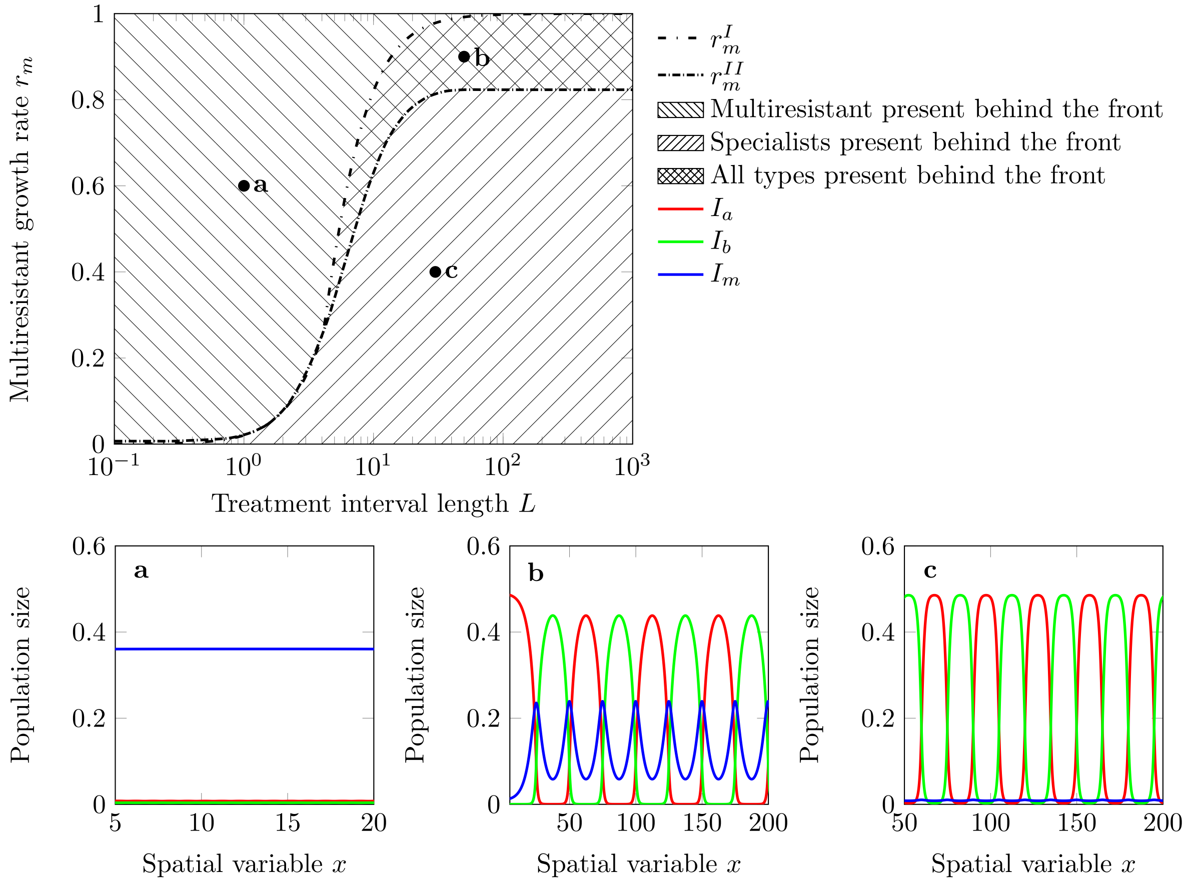

**Figure S6: Composition of the pathogen population far behind the front of the epidemic depending on coefficients $L$ and $r_m$.** The upper area (north-west hatches) corresponds to a stable steady-state where only the multiresistant type is present (as illustrated in (a)). The middle area (north-west and north-east hatches) corresponds to stable steady-states where all three types of pathogen are present (as illustrated in (b)). The lower area (north-east hatches) corresponds to stable steady-states where only the two specialists are present (as illustrated in (c)). Parameters: $\sigma = 1$, $\mu = 0$, $\beta_m = 1$, $r = 1$, $\mu = 10^{-3}$.

| **parameter** | **description** |
|---|---|
| $N$ | Number of hosts per deme |
| $\delta x$ | Distance between two successive demes |
| $\frac{\sigma}{2N\delta x^2}$ | Rate at which a couple of neighbouring hosts switch position |
| $\frac{\beta_a(i\delta x)}{N}$ | Rate a type $a$ infection affecting a given host is transmitted to a given susceptible host |
| $\frac{\beta_b(i\delta x)}{N}$ | Rate a type $b$ infection affecting a given host is transmitted to a given susceptible host |
| $\frac{\beta_m}{N}$ | Rate a type $m$ infection affecting a given host is transmitted to a given susceptible host |
| $\alpha$ | Recovery rate of an infected host |
| $\mu$ | Mutation rate of a pathogen of one type to a different type |

We can study the influence of demographic stochasticity with a change in the system size $N$. In particular, when $N \to \infty$ the population demographic stochasticity becomes vanishingly small (we refer to [7]for a related argument). It is convenient to track the densities of the different hosts instead of discrete number of hosts. If $S^N$ (resp. $I_a^N$, $I_b^N$, $I_m^N$) count the number of hosts, in each node, that are susceptible (resp. infected $a$, $b$, $m$),

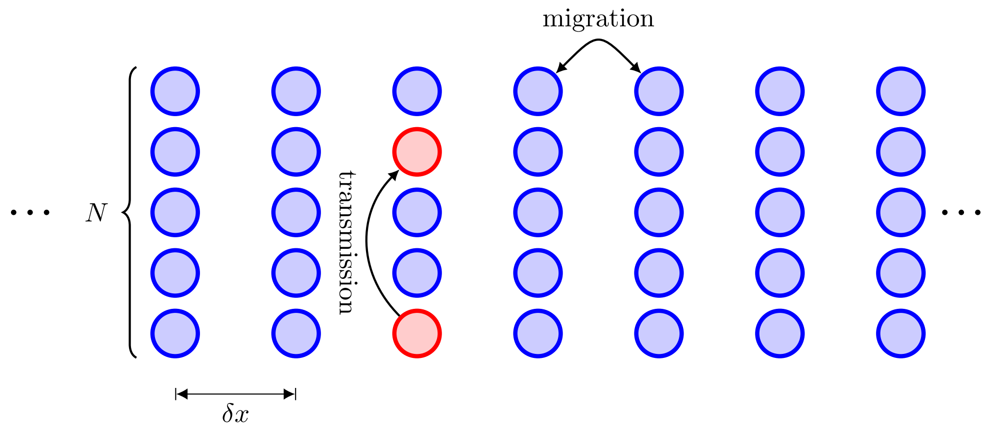

**Figure S7: Schematic representation of the lattice setting of the hosts.** Horizontal transmission occur within a site. Migration of the hosts occur between pairs of hosts at neighbour sites. If one of the neighboring host is infected this event leads to the migration of the pathogen.

then:

$$S := K\frac{S^N}{N}, \quad I_a := K\frac{I_a^N}{N}, \quad I_b := K\frac{{I_b}^N}{N}, \quad I_m := K\frac{{I_m}^N}{N}.$$

When $N \to \infty$ and $\delta_x \to 0$, this system converges to (2). In the following we compare the results of the deterministic model to the dynamics of this stochastic model with a finite number of hosts per deme.

#### 2.2 Propagation speed for simplified stochastic models

##### 2.2.1 Propagation speed of the multiresistant genotype

Just as for the full system, the stochastic model where only the multiresistant type is present converges to the corresponding deterministic model. As a consequence, when $N \gg 1$ and $0 < \delta x \ll 1$, we expect the propagation speed of the multiresistant type alone in the stochastic model to be close to $c_m = 2\sqrt{\sigma r_m}$. The impact of the population size on the propagation speed has been described in [4] and these results have been complemented in numerous subsequent manuscripts, see e.g. [2, 14]. When $N$ is large, the stochastic spread of the epidemic can then be approximated by:

$$\nu_m \sim 2\sqrt{\sigma r_m} - \frac{\kappa}{\left(\ln\left(\frac{K}{\delta x}\right)\right)^2}, \tag{14}$$

for some constant $\kappa > 0$. As illustrated by Figure S8a, this approximation provides a good qualitative description of the influence of $N$ on the propagation speed. For the present study, we will not use the precise approximation provided by (14), but simply remember that the deterministic model (4) does provide a satisfactory description of the speed of the propagation of the multiresistant type, at least when $N \geq 100$ (see Figure S8b).

##### 2.2.2 Propagation speed for a specialist population alone

When only one specialist type (e.g. type $a$) is present the pathogen population is spreading in a heterogeneous environment. Numerical simulations show that the speed $\nu_a$ for the stochastic model is always smaller than the speed $c_a$ of the deterministic model. We are not aware of rigorous mathematical work on the impact of the population size in this heterogeneous setting, but we expect that an approximation similar to (14) holds, that is:

$$\nu_a \sim c_a - \frac{\kappa}{\left(\ln\left(\frac{N}{\delta x}\right)\right)^2} \text{ when } N \gg 1, \tag{15}$$

for some constant $\kappa > 0$. We believe that this approximation will hold when $N$ is large compared to the spatial pattern length $L$. When $L$ is large compared to $N$, a different approximation should be used (see [15] for a related analysis). In practice, for any given parameter $N$, numerical simulations (see Figure S9) show that the propagatiopn speed $\nu_a$ becomes qualitatively different from its deterministic counterpart when $L$ is large.

Indeed, when $L$ is large, the specialist genotype is driven to extinction in the unfavorable habitat before it is able to reach the new favourable patch. In Section C, we show that if $1 \ll \ln\left(\frac{N}{\delta x}\right) \ll L$, the speed of the stochastic epidemic can be approximated by (see (21))

$$\nu_a \sim e^{-\frac{3L}{4}\sqrt{\frac{r}{\sigma}}+\ln(N/\delta x)}.$$

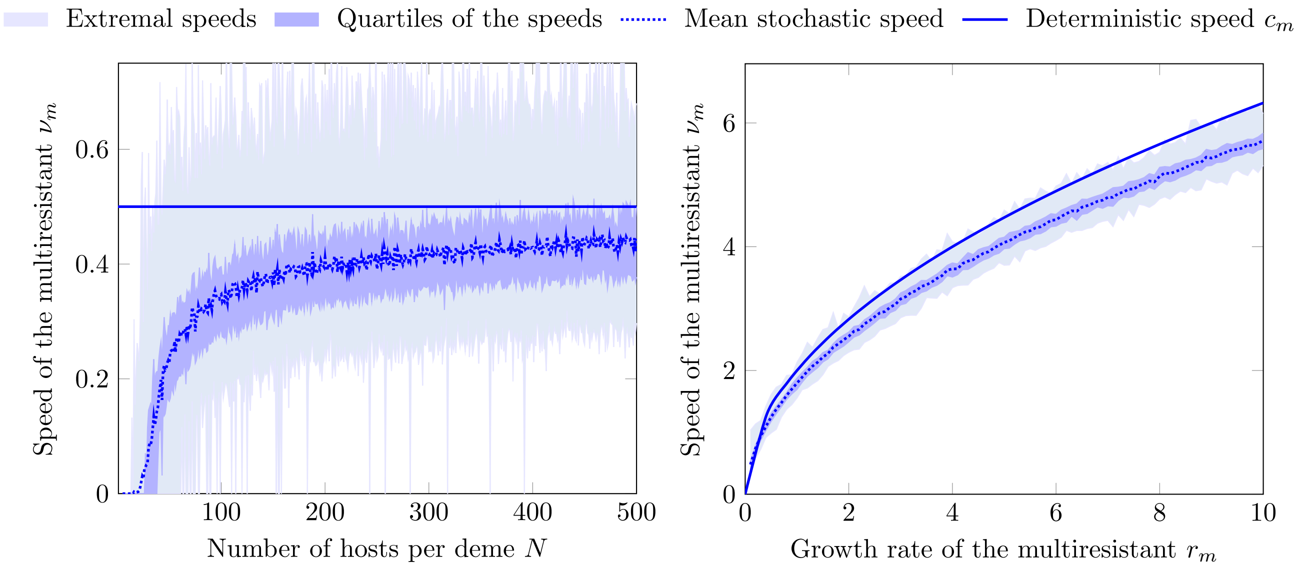

**(a)** $r_m = 1/16$.

**(b)** $N = 100$.

**Figure S8: Propagation speed of the generalist alone for the stochastic model.** We represent this speed as a function of the number of hosts per site $N$ with $\beta_m = 1.0 + 1/16$ fixed (first graph), and as a function of the growth rate of the multiresistant $r_m$ for $N = 100$ fixed (second graph). Other parameters: $\alpha = 1.0$, $\sigma = 1.0$, $\delta x = 0.1$, $\mu = 0$.

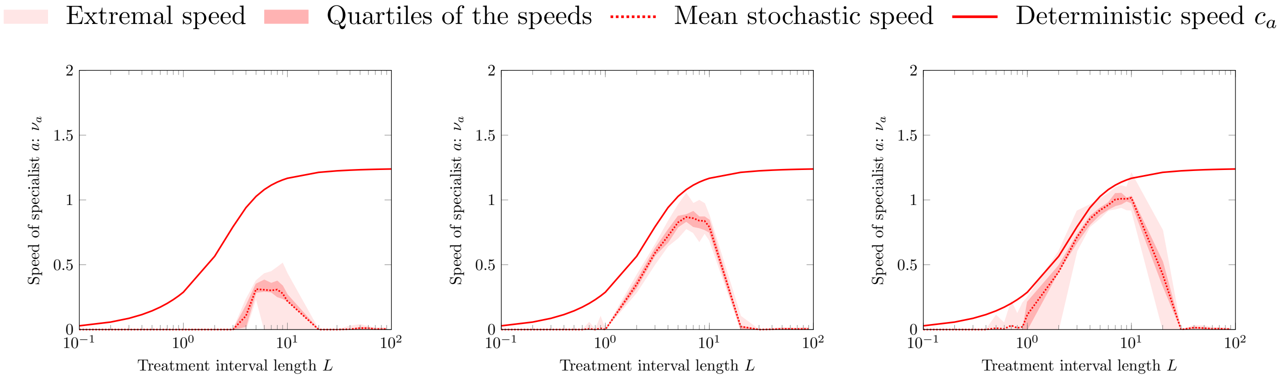

**(a)** $N = 10$

**(b)** $N = 100$

**(c)** $N = 1000$

**Figure S9: Impact of the local population size $N$ on the propagation speed of the stochastic model for a specialist alone**. Parameters: $\sigma = 1.0$, $\Delta x = 0.1$.

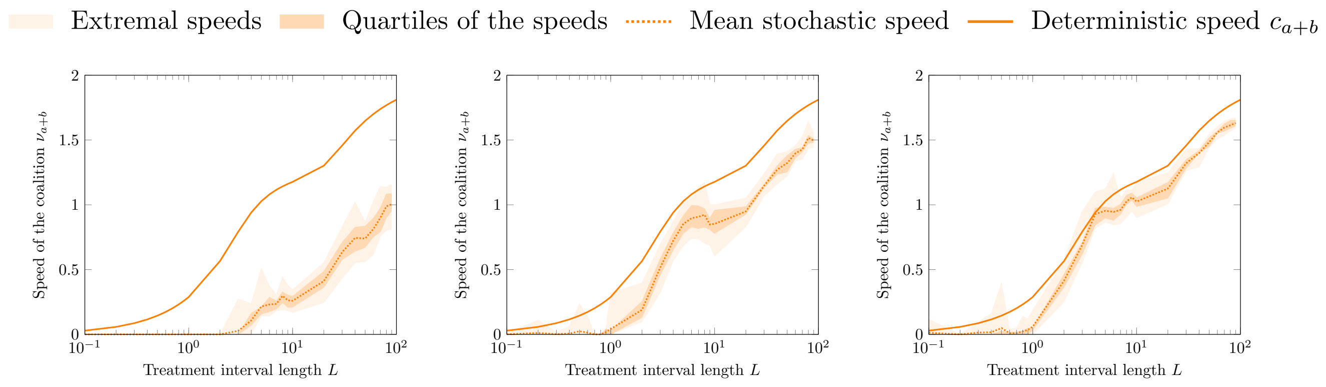

**(a)** $N = 10$. **(b)** $N = 100$. **(c)** $N = 1000$.

**Figure S10: Impact of the population size $N$ on the propagation speed of the stochastic model for a coalition of the two specialist types.** Parameters: $\alpha = 1.0$, $\beta = \{0.0, 2.0\}$, $\sigma = 1.0$, $\delta x = 0.1$, $\mu = 10^{-2}$ (coalition).

In the above expression, $L$ has an exponential impact on the speed while $N$ affects the speed linearly. The overwhelming effect of increasing $L$ captures the property illustrated by Figure S9, where the spread of the epidemic seems to come to a halt when $L$ goes beyond a threshold value

$$L \gg L_e := \frac{4}{3}\sqrt{\frac{\sigma}{r}} \ln\left(\frac{N}{\delta x}\right), \tag{16}$$

and this threshold value increase very slowly (ie. logarithmically) when $N$ increases.

##### 2.2.3 Propagation speed for a coalition of specialists

We now assume that only the two specialist genotypes $a$ and $b$ are present. Note that mutation between specialist genotypes allows the epidemic to spread even when $L$ becomes very large: the propagation speed of the coalition, illustrated by Figure S10 is positive for large $L$, which is not the case for a single specialist (see Figures S9). Just as in the deterministic case, the propagation speed of the coalition appears to be given by the *Independent propagation of specialists* if $L > 0$ is below a critical value $L_c \sim \frac{4}{1-3^{-3/4}\sqrt{2}}\sqrt{\frac{\sigma}{r}} \ln\left(\frac{r}{\mu}\sqrt{\frac{r}{\sigma}}\right)$, see (13). It is interesting to relate this quantity to the critical length $L_e = \frac{4}{3}\sqrt{\frac{\sigma}{r}} \ln\left(\frac{N}{\delta x}\right)$ (see (16)): if $L \geq L_e$, the stochastic effects bring a population composed of a single specialist type only to a halt (see Section 2.2.2). More broadly, the stochastic effect play an important role for a population of specialists alone if $L \gtrsim L_e$. The stochastic effects slowing down the propagation of a single specialist will then be reflected in the propagation of a coalition of specialists if $L_e \lesssim L_c$.

If $\mu > 0$ is fixed and $N > 0$ is sufficiently large, then $L_e \gg L_c$. The propagation speed of the coalition of specialists, $\nu_{a+b}$, is then well approximated by the speed derived for the deterministic model (Figure S10). For a given $N > 0$, if $\mu > 0$ is small enough, $L_e \lesssim L_c$ and the stochastic effects on the propagation speed of a specialist alone are present for intermediate lengths $L$, as can be seen in Fig. S11: for $L \leq 20$, the propagation speed of the coalition is well approximated by the non-monotonic speed of a single specialist type as given represented in Fig. S9.

As a consequence, the speed of the epidemic can be limited by the amount of mutation if $L$ is in an intermediate parameter range ($L \in [10, 100]$ in Figure S11), and this creates a drop in the propagation speed for intermediate parameters $L$ that was not present in the deterministic case.

#### 2.3 Analysis of the full stochastic model

As illustrated by Figure S10, the propagation speed of the full stochastic model is well approximated by the deterministic model if the number of hosts per site $N$ is large. The stochasticity however slows down the propagation for any heterogeneity parameter $L$.

There is a more pronounced slow down effect if $L$ is in an intermediate range and if $\mu > 0$ is small: this can be observed in Figure S13 for $L \in [10, 100]$, and it originates from the stochastic effects on the propagation of a coalition of specialists that has been described in Section 2.2.3. A consequence of this effect is that the multiresistant pathogens can become the one that propagates the fastest for these intermediate parameters $L$. This means the multiresistant pathogen can appear on the edge of the propagation front for a larger set of parameters: this can be seen in Figure 6, where the blue area extends further down for $L \in [10, 100]$ than in its deterministic counterpart.

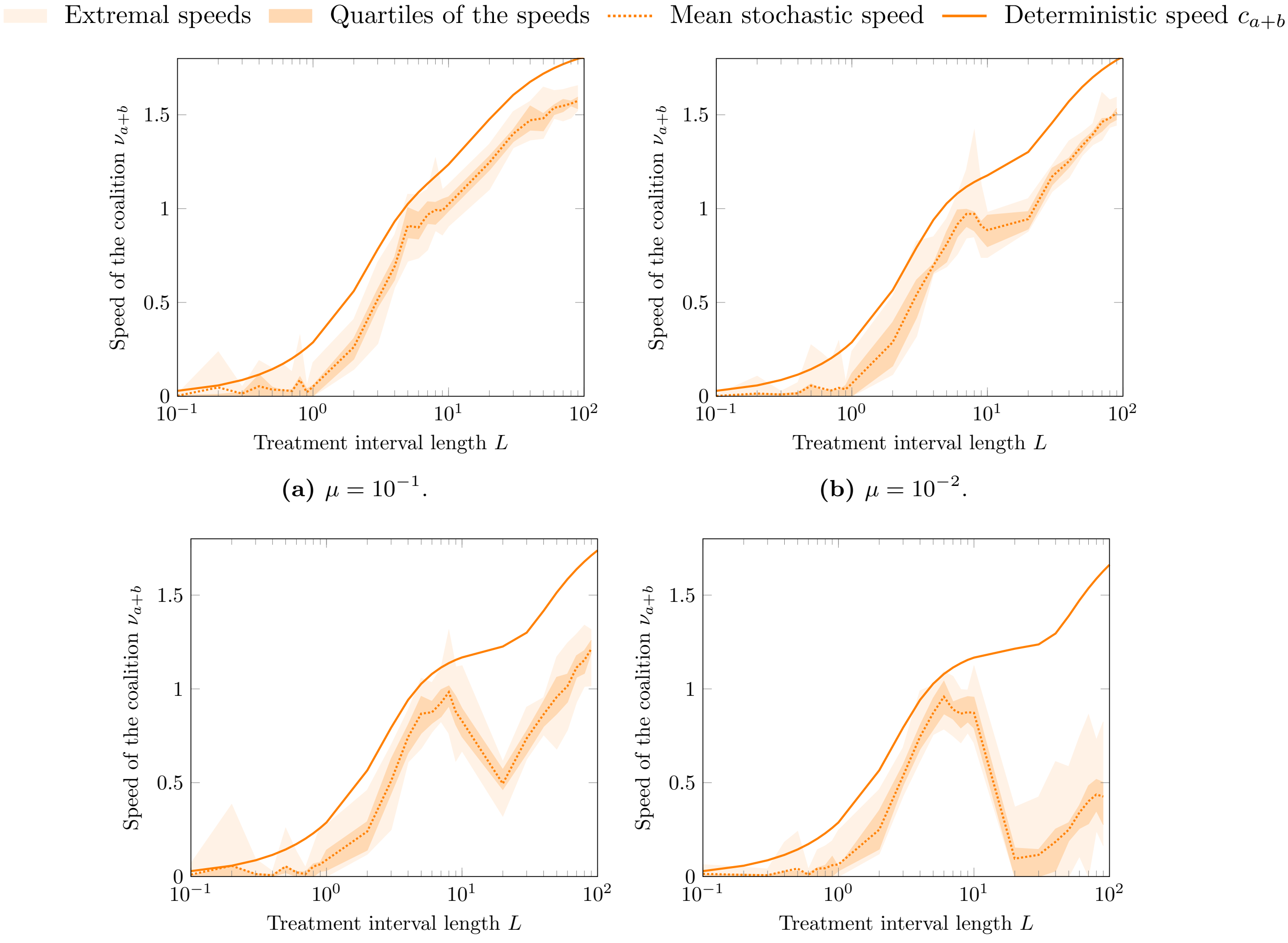

**(a)** $\mu = 10^{-1}$.

**(b)** $\mu = 10^{-2}$.

**(c)** $\mu = 10^{-3}$.

**(d)** $\mu = 10^{-4}$.

**Figure S11: Impact of the mutation rate $\mu$ on the propagation speed of the stochastic model for a coalition of the two specialists.** Parameters: $\alpha = 1.0$, $\beta = \{0.0, 2.0\}$, $\sigma = 1.0$, $\Delta x = 0.1$, $N = 100$.

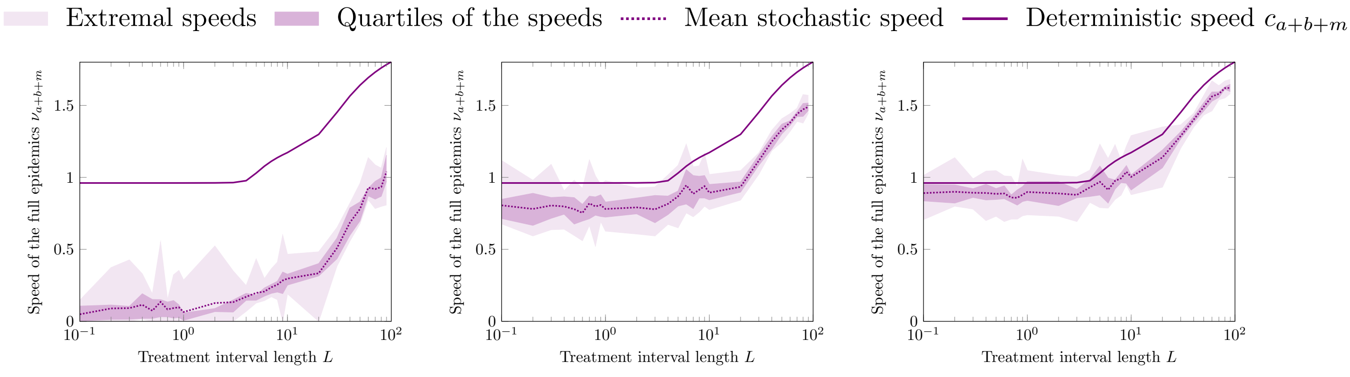

**(a)** $N = 10$.

**(b)** $N = 100$.

**(c)** $N = 1000$.

**Figure S12: Impact of the population size $N$ on the propagation speed of the full model.** Parameters: $\alpha = 1.0$, $\beta = \{0.0, 2.0\}$, $\sigma = 1.0$, $\delta x = 0.1$, $\beta_m = 1.0 + 1/16$, $\mu = 10^{-2}$.

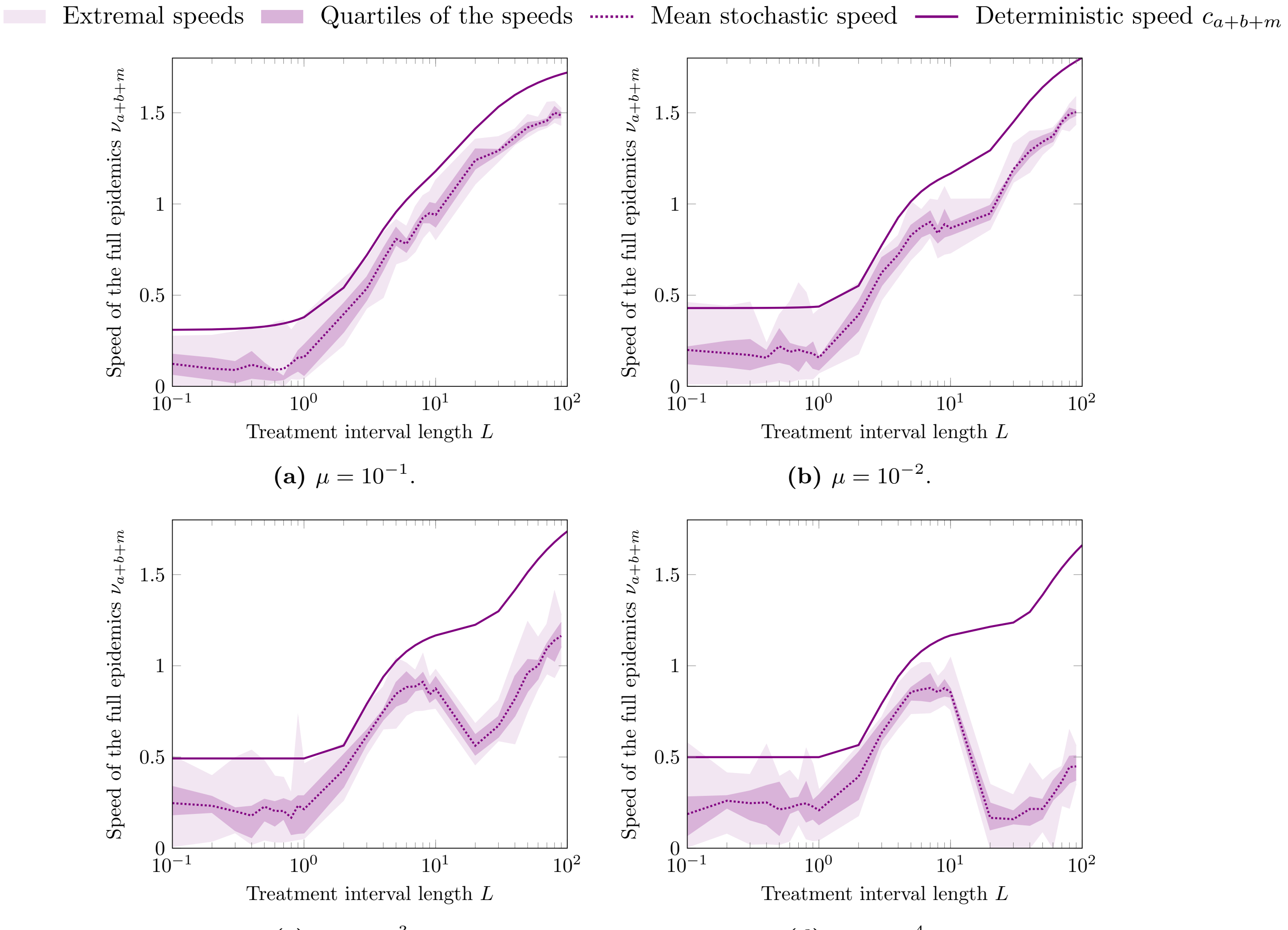

**(a)** $\mu = 10^{-1}$.

**(b)** $\mu = 10^{-2}$.

**(c)** $\mu = 10^{-3}$.

**(d)** $\mu = 10^{-4}$.

**Figure S13: Impact of the mutation rate $\mu$ on the propagation speed of the full stochastic model.** Parameters : $\alpha = 1.0$, $\beta = \{0.0, 2.0\}$, $\sigma = 1.0$, $\Delta x = 0.1$, $\beta_m = 1.0 + 1/16$, $N = 100$.

### Appendix

### A Remarks on the analysis method

In our analysis, we have considered the speed of sub-units of the population model: We considered the types separately, and the two specialist types without the multiresistant type. This turned out to be an efficient analysis method for both deterministic (see Section 1.2.1 and 1.2.2) and stochastic models (see Section 2.2.3, 2.3).

This idea of the propagation speed being the maximum of the propagation speeds of sub-units of the system has already been used in [10], where the model was a system of reaction-diffusion equation and the environment was constant. The propagation speed of the population can then be related to the principal eigenvalue of the linearized system on $[-R,R]$ (for $R>0$ arbitrarily large):

$$\mathcal{L}((u_i)_i) = \begin{cases} \sigma\Delta_x u_1 + r_1 u_1 - \left(\sum_{j\neq 1}\mu_{1j}\right)u_1 + \sum_{j\neq 1}\mu_{j1}u_j, \\ \quad\vdots \\ \sigma\Delta_x u_n r_n u_n - \left(\sum_{j\neq n}\mu_{nj}\right)u_n + \sum_{j\neq n}\mu_{jn}u_j, \end{cases}$$

while $(u_i)_i(\pm R)=0$. The principal eigenvalue of this system is the largest $\lambda\in\mathbb{R}$ such that for some $(u_i)_i\neq 0$, $\mathcal{L}((u_i)_i)=\lambda(u_i)_i$ and $(u_i)_i(\pm R)=0$. One may show that this is achieved with $\lambda\sim_{R\gg 1}\bar\lambda_\mu$, $(u_i)_i(x)=\sin\left(\frac{\pi}{2R}(x+R)\right)(\Gamma_i)_i$, where $(\bar\lambda_\mu,(\Gamma_i)_i)$ are the principal eigenvalue and eigenvector of the matrix

$$A_\mu = \begin{pmatrix} r_1-\left(\sum_{j\neq 1}\mu_{1j}\right) & \mu_{21} & \mu_{31} & \cdots & \mu_{(n-1)1} & \mu_{n1} \\ \mu_{21} & r_2-\left(\sum_{j\neq 2}\mu_{2j}\right) & \mu_{32} & \cdots & \mu_{(n-1)2} & \mu_{n2} \\ & & \ddots & & & \\ & & & \ddots & & \\ \mu_{1(n-1)} & \mu_{2(n-1)} & \mu_{3(n-1)} & \cdots & r_{n-1}-\left(\sum_{j\neq n-1}\mu_{(n-1)j}\right) & \mu_{n(n-1)} \\ \mu_{1n} & \mu_{2n} & \mu_{3n} & \cdots & \mu_{(n-1)n} & r_n-\left(\sum_{j\neq n}\mu_{nj}\right) \end{pmatrix}.$$

Since $A_\mu=\mathrm{diag}((r_i)_i)+\mathcal{O}((\mu_{ij}))$, the principal eigenvalue of $A_\mu$ satisfies $\lambda_\mu=\max_i r_i+\mathcal{O}((\mu_{ij}))$. The propagation speed of the population thus satisfies

$$c_\mu = 2\sqrt{\sigma\max_i r_i}+\mathcal{O}((\mu_{ij})).$$

In this study, we extend this idea along two axes: the case of a PDE model in a heterogeneous (periodic) environment and a stochastic model. The simple argument presented above cannot easily be extended to these more complicated settings. We do not have a rigorous argument for these extensions, and it would be really interesting to investigate this idea in future works. We describe below an argumentation frame that may be attempted to reach a better understanding of these extensions.

The propagation models we have worked with in this study are of *pulled* type: their propagation speed is guided by individuals far ahead of the bulk of the population. This property implies that the propagation is guided by unusual events, that is individuals migrating far ahead of the front and/or individuals that have an unusually large progeny. Since these events are rare, there will be enough time between two such events so that "almost every possible strategy" will be attempted in the mean time. As a consequence, the speed of the population can be computed as the maximum of the speeds corresponding to each possible strategy. The mathematical setting to describe the importance of the strategies maximizing the propagation speed is likely to be large deviation theory (see e.g. [22]). This theory has a deterministic alter-ego: the theory of Hamilton-Jacobi equations, that are obtained from reaction-diffusion equation through a WKB transformation. This theory provides a fruitful connection between probability theory, PDE and optimization and it could provide an interesting point of view in this context also.

### B Numerical estimates of the speed of the deterministic front

To estimate the propagation speed of the whole system, a first idea is to compute a mean of the displacement over a long period of time. While it is worth implementing this algorithm and using it to some extent to compare it with other methods, it suffers from several limitations:

- A choice need to be made on the numerical method to cope with an unbounded space (very large grid? Shifting box? If so, boundary conditions?)
- Which characterization should be used to measure the progression of the front: small-value level set at the edge of the front? How is this impacted by the discretization method?
- Computational time: if a large spatial interval is considered, the computing time becomes too large.

Luckily, in our case, we are able to compute this speed through an eigenvalue problem.

**The eigenvalue problem** We chose to use the implicit formulation for the speed of the front:

$$c = \inf_{\lambda>0} -\frac{k(\lambda)}{\lambda}, \tag{17}$$

where $k(\lambda)$ is the principal eigenvalue of the associated linearized problem:

$$\begin{cases} -\varphi_{xx} + 2\lambda\varphi_x - \lambda^2\varphi - A(x)\varphi = k(\lambda)\varphi, \qquad x \in \mathbb{R} \\ \varphi \text{ is } L-\text{periodic}, \quad \varphi > 0, \end{cases} \tag{18}$$

where the matrix $A(x)$ encodes the reaction terms specific to the problem.

This formulation has the advantage of linking the computation of the speed of the problem with two classical problems in numerical analysis: optimizing a real function and computing an eigenvalue of a linear problem. One other advantage of this approach is that the computed speed is the exact travelling wave speed for propagation model in the discrete lattice, thanks to the results of Weinberger [18].

There are numerous ways to approximate the principal eigenvalue in (18). Among those, the *power method* is particularly interesting since it is simple to implement and it allows us to directly compute the principal eigenvalue of a matrix with no requirement of symmetry (Problem (18) is not self-adjoint when $\lambda \neq 0$). Moreover, if well implemented, it preserves the positivity of the initial vector, which is a requirement to find the principal eigenvalue of the limit problem.

**Discretization and numerical scheme** We discretize the periodicity cell $[0, L]$ in $N$ points $(x_i)_{i\in\{1,\dots,N\}} = (\Delta x, 2\Delta x, \dots, L)$ with $\Delta x = \frac{L}{N}$. We denote $\varphi_i = \varphi(x_i)$. We use the following approximations for the derivatives of $\varphi$:

$$\begin{aligned} \partial_x\varphi(x_i) &= \frac{\varphi_{i+1} - \varphi_i}{\Delta x} + \mathcal{O}(\Delta x), && i \in \{1, \dots, N-1\} \\ \partial_x\varphi(x_N) &= \frac{\varphi_1 - \varphi_N}{\Delta x} + \mathcal{O}(\Delta x), \\ \partial_{xx}\varphi(x_i) &= \frac{\varphi_{i-1} - 2\varphi_i + \varphi_{i+1}}{\Delta x^2} + \mathcal{O}(\Delta x^2), && i \in \{2, \dots, N-1\} \end{aligned}$$

thus reflecting the periodic boundary conditions $\varphi(0) = \varphi(L)$. We denote $D_N$ the matrix approximating the left-hand side of (18) by this mean. For 1 species, the discrete approximation of the left-hand side of (18) writes as the matrix

$$D_N = \begin{pmatrix} a_1 & b_1 & 0 & \cdots & 0 & c_n \\ c_1 & a_2 & b_2 & \cdots & & 0 \\ 0 & c_2 & a_3 & & & \vdots \\ \vdots & & & \ddots & & \\ 0 & & & & a_{n-1} & b_{n-1} \\ b_n & 0 & \cdots & & c_{n-1} & a_n \end{pmatrix} \tag{19}$$

with $a_i = \frac{2\sigma}{\Delta x^2} + 2\frac{\lambda}{\Delta x} - r(x_i) - \lambda^2$, $b_i = -\frac{\sigma}{\Delta x^2} - 2\frac{\lambda}{\Delta x}$, and $c_i = -\frac{\sigma}{\Delta x^2}$. For two or more species, the matrix $D_N$ is still sparse but much more difficult to represent because of the mutation terms.

Our goal is to find the principal eigenpair $(k_N, \phi_N)$ of $D_N$ as an approximation for the principal eigenpair $(k(\lambda), \varphi)$ solution to (18). To this end we investigate the slightly modified problem:

$$\tilde{D}_N\phi_N = \mu_N\phi_N, \tag{20}$$

where $\tilde{D}_N = D_N + (\max_{x\in\mathbb{R}} r(x) + \lambda^2)I_N$. We use the following result on the modified matrix $\tilde{D}_N$, which are classical.

**Proposition 1** (Positivity)**.** *The matrix $\tilde{D}_N$ has the following properties:*

1. *if $Y \geq 0$ and $\tilde{D}_N X = Y$, then $X \geq 0$.*

2. *if $Y \geq 0$, $Y \not\equiv 0$ and $\tilde{D}_N X = Y$, then $X > 0$.*

We also use the following lemma, that is a consequence of the first lemma and the Perron-Frobenius Theorem applied to $\tilde{D}_N^{-1}$:

**Proposition 2.** *There exists a unique eigenpair $(\mu_N, \phi_N > 0)$ solution to (20).*

We define the approximating sequence $(X_n)_{n\in\mathbb{N}}$, where $X_0$ is a given nonnegative nontrivial vector and $X_{n+1} = \frac{\tilde{D}_N^{-1} X_n}{\|\tilde{D}_N^{-1} X_n\|}$.

**Proposition 3** (Inverse power method)**.** *The sequence $X_n$ converges exponentially fast to $\phi_N$.*

Once a good approximation of $\phi_N$ is known, we easily retrieve $k_N$, and thus the propagation speed for the discretized problem (see (17)). We can estimate numerically the errors in the case of two species. As expected, the method appears to be of order 1 in $\Delta x$.

### C Probability of an individual crossing of an unfavourable zone

In this section, we estimate the probability that an individual succeeds to cross a stretch of unfavourable environment of length $L/2$, when $L$ is large. This is equivalent to the probability that an individual starting at the origin ($x = 0$) reaches a position $x \geq \frac{L}{2}$ in a homogeneous unfavourable environment. In this unfavourable environment, there is no reproduction, individuals die at rate $r > 0$ and disperse at rate $\frac{\sigma}{\delta x^2}$, with a balanced dispersion distance of $\pm\delta x$.

**Remark 1.** The assumption that individuals do not reproduce will simplify our analysis. Considering situations where reproduction events are present even in the unfavourable zones would however be possible, using moment methods and many-to-few lemma.

Let $\bar{X}_i$ independent uniformly distributed random variables with

$$\bar{X}_i = \begin{cases} 1 \text{ with probability } \frac{1}{2}, \\ -1 \text{ with probability } \frac{1}{2}. \end{cases}$$

Following the notations from *Large deviations Theory*, we define $\Lambda_{\bar{X}_1}$ and the rate function $I_{\bar{X}_1}$ associated to dispersion events as follows:

$$\Lambda_{\bar{X}_1}(\lambda) := \ln\left(\mathbb{E}\left(e^{\lambda \bar{X}_1}\right)\right),$$

$$I_{\bar{X}_1}(x) := \sup_{\lambda\in\mathbb{R}} \left(\lambda x - \Lambda_{\bar{X}_1}(\lambda)\right).$$

The rate function $I_{\bar{X}_1}$ can be explicitly computed through the distribution function of $\sum_{i=1}^n \bar{X}_i$:

$$I_{\bar{X}_1}(x) = \frac{(1+x)\ln(1+x) + (1-x)\ln(1-x)}{2}.$$

The process $(\bar{X}_i)_i$ satisfies a Large Deviation Principle, and then, thanks to the Cramér's Theorem [22], for any $\delta x > 0$,

$$\frac{1}{n}\ln \mathbb{P}\left(\frac{\sum_{i=1}^n X_i}{n} > x\right) = \frac{1}{n}\ln \mathbb{P}\left(\frac{\sum_{i=1}^n \bar{X}_i}{n} > \frac{x}{\delta x}\right) \to_{n\to\infty} -I_{\bar{X}_1}\left(\frac{x}{\delta x}\right).$$

Let $Y(t)$ be the position of the individual we consider, and define $\bar{Y}(t) := \sum_{i\in J(t)} X_i$, where $J(t)$ is the cumulative distribution of the dispersion events affecting the individual (i.e. a Poisson law of parameter $\frac{\sigma}{\delta x^2}$). Let denote by $T$ the stopping time defining the death of the individual. For $t \leq T$, $Y(t)$ and $\bar{Y}(t)\delta x$ have the same law. We have, thanks to the independence between death events and dispersion events,

$$\begin{aligned}\mathbb{P}(\exists t \leq T; Y(t) > L/2) &= \int_0^\infty \mathbb{P}(\max_{\sigma\in[0,s]} Y(\sigma) > L/2 | T = s)\, d\mathbb{P}(T = s)(s) \\ &= \int_0^\infty \mathbb{P}\left(\max_{\sigma\in[0,s]} \bar{Y}(\sigma) > \frac{L}{2\delta x}\right) d\mathbb{P}(T = s)(s).\end{aligned}$$

We use next the reflection principle, which is valid when $\delta x \ll L$: $\delta x\,\bar{Y}(t)$ is then very close to a Brownian motion.

$$\mathbb{P}(\exists t \le T;\, Y(t) > L/2) \sim_{\delta x \ll 1} 2\int_0^\infty \mathbb{P}\left(\bar{Y}(s) > \frac{L}{2\delta x}\right) d\mathbb{P}(T = s)(s),$$

and we proceed to estimate this quantity

$$\begin{aligned}
&\mathbb{P}(\exists t \le T;\, Y(t) > L/2)\\
&\quad \sim 2\int_0^\infty \left(\sum_{n=0}^\infty \mathbb{P}\left(\frac{\sum_{i=1}^n \bar{X}_i}{n} > \frac{L}{2\delta x}\right)\mathbb{P}\left(n \text{ dispersion events in } [0,s]\right)\right) d\mathbb{P}(T = s)(s)\\
&\quad = 2\int_0^\infty \sum_{n=0}^\infty \mathbb{P}\left(\frac{\sum_{i=1}^n \bar{X}_i}{n} > \frac{L}{2\delta x}\right)\left(\frac{\sigma}{\delta x^2}s\right)^n \frac{e^{-\frac{\sigma}{\delta x^2}s}}{n!} r e^{-rs}\, ds\\
&\quad \sim 2\int_0^\infty \sum_{n=0}^\infty e^{-nI_{\bar{X}_1}\left(\frac{L}{2n\delta x}\right)}\left(\frac{\sigma}{\delta x^2}s\right)^n \frac{e^{-\frac{\sigma}{\delta x^2}s}}{n!} r e^{-rs}\, ds\\
&\quad = 2\sum_{n=0}^\infty r e^{-nI_{\bar{X}_1}\left(\frac{L}{2n\delta x}\right)}\frac{\sigma^n}{\delta x^{2n} n!}\int_0^\infty s^n e^{-\left(\frac{\sigma}{\delta x^2}+r\right)s}\, ds = 2\sum_{n=0}^\infty r e^{-nI_{\bar{X}_1}\left(\frac{L}{2n\delta x}\right)}\frac{\sigma^n}{\delta x^{2n} n!}\frac{n!}{(\sigma/\delta x^2 + r)^{n+1}}\\
&\quad = \frac{2r}{\sigma/\delta x^2 + r}\sum_{n=0}^\infty e^{-n\left(I_{\bar{X}_1}\left(\frac{L}{2n\delta x}\right)+\ln\left(1+\frac{r}{\sigma}\delta x^2\right)\right)} \sim \frac{2r}{\sigma/\delta x^2 + r} e^{-L\inf_{x>0}\frac{I_{\bar{X}_1}\left(\frac{x}{2\delta x}\right)+\ln\left(1+\frac{r}{\sigma}\delta x^2\right)}{x}}.
\end{aligned}$$

The infimum is reached for $\bar{x} > 0$ such that $\frac{1}{2\bar{x}\,\delta x} I'_{\bar{X}_1}\left(\frac{\bar{x}}{2\delta x}\right) = \frac{I'_{\bar{X}_1}\left(\frac{\bar{x}}{2\delta x}\right)+\ln\left(1+\frac{r}{\sigma}\delta x^2\right)}{\bar{x}^2}$. Since $I'_{\bar{X}_1}(0) = 0$ and $I''_{\bar{X}_1}(0) = \mathrm{Var}(\bar{X}_1) = 1$,

$$\bar{x} \sim_{\delta x \ll 1} 2\sqrt{\frac{r}{\sigma}}\delta x^2,$$

and thus

$$\mathbb{P}(Y(t) > L/2) \sim e^{-\frac{3L}{4}\sqrt{\frac{r}{\sigma}}}$$

The probability, given that the population is present in the favourable zone $[nL, (n+1/2)L]$, that an individual situated close to $(n+1/2)L$ at time $t$ succeeds to reach the next favourable zone $[(n+1)L, (n+3/2)L]$ is then of the order of $\frac{N}{\delta x}e^{-\frac{3L}{4}\sqrt{\frac{r}{\sigma}}}$. Successful crossing then happen at a rate of the order of $e^{-\frac{3L}{4}\sqrt{\frac{r}{\sigma}}+\ln(N/\delta x)}$. As soon as $L \gg \ln(N/\delta x)$, waiting for successful crossings of unfavourable zones has a dominant effect on the propagation speed of the population that we can thus estimate: If $L \gg \ln(N/\delta x)$, the propagation speed of a single population of specialists is of the order of

$$\nu_a \sim e^{-\frac{3L}{4}\sqrt{\frac{r}{\sigma}}+\ln(N/\delta x)} \quad \text{if } L \gg 1. \tag{21}$$

[22] Ofer Zeitouni and Amir Dembo, *Large deviations techniques and applications*, 1998.
